# Central place foragers, prey depletion halos, and how behavioral niche partitioning promotes consumer coexistence

**DOI:** 10.1101/2024.06.13.598783

**Authors:** Claus Rueffler, Laurent Lehmann

## Abstract

Many seabirds congregate in large colonies for breeding, a time when they are central place foragers. An influential idea in seabird ecology posits that competition during breeding results in an area of reduced prey availability around colonies, a phenomenon known as Ashmole’s halo, and that this limits colony size. This idea has gained empirical support, including the finding that species coexisting within a colony might be able to do so by foraging on a single prey species but at different distances. Here, we provide a comprehensive mathematical model for central place foragers exploiting a single prey in a two-dimensional environment, where the prey distribution is the result of intrinsic birth and death, movement in space and mortality due to foraging birds (we also consider a variant tailored toward colonial social insects). Bird predation at different distances occurs according to an ideal free foraging distribution that maximizes prey delivery under flight and search costs. We fully characterize the birds’ ideal free distribution and the prey distribution it generates. Our results show that prey depletion halos around breeding colonies are a robust phenomenon and that the birds’ ideal free distribution is sensitive to prey movement. Furthermore, coexistence of several seabird species on a single prey easily emerges through behavioral niche partitioning whenever trait differences between species entail trade-offs between efficiently exploiting a scarce prey close to the colony and a more abundant prey far away. Such behavioral-based coexistence-inducing mechanism should generalize to other habitat and diet choice scenarios.

**Significance statement:** This study presents a mathematical model to explore the distribution of foraging trips among seabirds breeding on isolated islands, providing insight into the emergence of prey depletion halos around colonies. Our findings reveal that such halos are a robust feature of central place foraging, independent of prey dynamics. Additionally, the model shows that trait-mediated niche partitioning promotes coexistence among species through behavioral segregation into different circular zones around the island. This partitioning occurs despite a shared preference to forage close to the island, where flight costs are lowest. The study advances understanding of ecological coexistence mechanisms and suggests broader applicability to other predator-prey systems beyond seabird ecology, offering a new perspective on community assembly under shared preferences.

## 1 Introduction

Seabirds breed in very large colonies (Furness and Monaghan, 1987; Coulson, 2002; Mitchell et al., 2004; Patterson et al., 2022), and various mechanisms have been proposed to explain the limits to colony size. The most influential one is due to Ashmole (1963), who hypothesized that colony size is regulated through negative density-dependence acting during the breeding season. He reasoned that during breeding, seabirds only forage in a restricted neighborhood of the colony because time intervals between visits to the nest cannot be too long and energetic costs for foraging trips cannot be too high if prey is to be delivered at a sufficiently high rate to nestlings. This area-restricted foraging process should lead to a zone of prey depletion around large colonies, a phenomenon later dubbed *Ashmole’s halo* (Birt et al., 1987), *and this depletion* ultimately limits colony size.

While Ashmole’s hypothesis has received strong empirical support (Furness and Birkhead, 1984; Birt et al., 1987; Lewis et al., 2001; Forero et al., 2002; Ainley et al., 2003; Elliott et al., 2009; Oppel et al., 2015; Weber et al., 2021), to date no mathematical model exists that analyzes how individual behavioral decisions of central place foragers about their foraging trips, the demography of these central place foragers and of their prey, jointly determine colony size and prey depletion halos. Existing models of central place foragers incorporate some of these aspects while neglecting others. Assuming that foragers distribute their foraging trips according to the ideal free distribution in a two-dimensional environment, Dukas and Edelstein-Keshet (1998) derive the shape of this distribution, but without incorporating prey dynamics. Thus, a corresponding prey depletion halo is not characterized. Gaston et al. (2007), again assuming that foragers adopt an ideal free distribution, derive the shape of prey depletion halos as they result from the energy requirements of different seabirds. However, the shape of the ideal free distribution is not characterized, precluding an understanding of the interdependence of predator behavior and prey depletion patterns. Lastly, Weber et al. (2021) study the emergence of a prey depletion halo given an empirically observed forager distribution, which again does not inform us about the interdependence of foragers and the distribution of their prey distributions. Formalizing a model for central place foragers based on an explicit description of prey population dynamics, the behavioral choice of predators about their foraging grounds, and their demography is necessary to determine the robustness and shape of prey depletion halos. Additionally, this allows us to understand how these halos are coupled to the forager’s ideal free distribution and how both prey depletion halos and foraging distributions jointly determine the forager’s equilibrium colony size.

For seabirds, the interaction between central place foragers and prey depletion halos is even more complex, since it is common for colonies to consist of several different species. This raises the question of what allows these species to coexist (Petalas et al., 2024), and how prey depletion halos form from their joint foraging effort. A study by Weber et al. (2021), conducted on Ascension Island, the place of Ashmole’s original research, found that different species of seabirds preferentially forage at different distances from the breeding colony on Ascension Island. Brown boobies (Sula leucogaster) forage closest to the island, masked boobies (*Sula dactylatra*) at intermediate distances, and Ascension frigatebirds (*Fregata aquila*) the furthest away. Weber et al. (2021) also provide evidence that these species all feed on flying fish, primarily tropical two-winged flying fish (*Exocoetus volitans*).

These findings suggest that different bird species can coexist by partitioning foraging space, even if this space is inhabited by only a single prey species. This idea is somewhat perplexing since the area close to the colony should be preferred by all species as foraging therein comes at the lowest costs, both in terms of time and energy expenditure. This means that central place foragers coexisting on a single prey species form a community with shared preferences (Rosenzweig, 1991). To understand under what conditions a pattern as observed by Weber et al. (2021) can indeed result from the joint interaction of several predator species, each adopting the ideal free distribution, and a single prey species, is sufficiently complex to again require a mathematical model. Such a formal analysis seems particularly interesting, given that the bulk of classical ecological coexistence theory is premised on individuals having distinct preferences, where species are optimal and prefer a different part of niche space (MacArthur and Levins 1967; MacArthur 1972; see Case 2000; Mittelbach and McGill 2019; Begon and Townsend 2021 for textbook treatments).

We here present a comprehensive mathematical model describing a population of central place foragers feeding on single prey species in a two-dimensional environment. Our goal is to understand how individual behavioral foraging decisions together with predator and prey dynamics jointly determine the forager’s ideal free distribution, its equilibrium population size and the prey’s equilibrium distribution in the vicinity of the colony (a variant of the model tailored toward colonial social insects is considered in Box 1). In the full model, we allow for an arbitrary number of predator species that differ in their foraging traits under the assumption that these cannot be optimized simultaneously but are coupled by trade-offs. For instance, in seabirds, species that are efficient flyers, and thereby incur low energetic costs from long distances foraging trips, might be poor at catching fish and vice versa. We aim at characterizing the conditions under which different species can coexist, in particular, whether coexistence is achieved by species partitioning the foraging area around the colony despite having a shared preference for foraging close to the colony. Our model is strongly inspired by the biology of seabirds breeding in large colonies on isolated island, and we therefore cast our model in terms of birds and fish, though our model is not meant to be an accurate description of any particular seabird population.

## 2 Model

### 2.1 Biological assumptions

We consider a population of birds breeding on an isolated island. The number of successfully raised offspring depends on the amount of prey they catch by foraging in the surrounding ocean and that they are able to deliver back to their nestlings. The waters surrounding the island are assumed to be homogeneous and harbor a single prey species. Any two points at equal distance *z* from the island are assumed to be visited with equal probability by foraging birds. As a result, the distribution of prey is radially-symmetric around the island. We denote by *R*(*z*) the equilibrium prey density at distance *z* from the island.

A foraging trip starts with birds flying from the island in a random direction to a point at distance *z*, where they start foraging. Individuals forage at a given location until they catch a prey, that is, individuals are single prey loaders (we consider multiple prey loaders in Appendix A.1). After a successful catch, birds fly back to the island to deliver the prey to their nest. If birds fly with speed *v*, then it takes a flying time of *T*_f_(*z*) = 2*z*/*v* time units to travel both directions.

The rate at which birds foraging at a point at distance *z* catch prey is *aR*(*z*), where *a* denotes the capture efficiency. The time it takes a bird to successfully catch a prey item at distance *z* is then exponentially distributed with mean *T*_s_(*R*(*z*)) = 1/(*aR*(*z*)), referred to as search time. If birds can extract on average *b* energy units from a single prey and if *c*_f_ and *c*_s_ denote the energy costs per time unit flying and searching, respectively, then

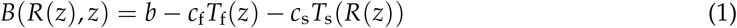

gives the net energy content of a prey item from distance *z*. In Appendix A.1, we show that with the above assumptions the rate at which birds deliver prey to the nest is given by

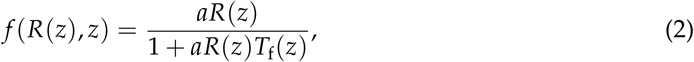

which we recognize as Holling’s type-II functional response (Holling, 1959). The net rate of energy delivery resulting from foraging at distance *z*, referred to as *payoff* and denoted Π(*z*), equals the product of the prey delivery rate and the net energy content per prey, and thus is

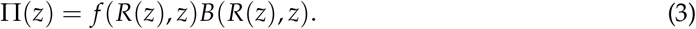

Hence, payoff at distance *z* depends on prey density at that distance, both through the functional response and the net energy content.

We assume that there are no prey on the island (i.e., *R*(0) = 0). Away from the island, equilibrium prey density *R*(*z*) at distance *z* depends on three processes. First, prey regrows at rate *g*(*R*(*z*)), given by the difference between birth and death that cannot be attributed to predation by birds breeding on the island. Second, prey move homogeneously in space at rate *m* (formally, we assume a reaction-diffusion process for the prey, see Appendix A.2 for details). Third, prey density is reduced due to mortality from foraging birds. Death through predation at distance *z* depends on how often an area at that distance is visited by birds, which, in turn, depends on the total number of birds and the probability that birds visit an area at distance *z*. In the absence of bird predation, prey have the same equilibrium density *R*^*^ everywhere around the island.

The probability of birds foraging at a certain distance is assumed to be an individual decision variable, which thus reflects the birds’ choice. We denote by *p*(*z*) the probability density that an individual forages at distance 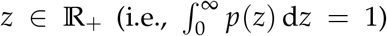 at a behavioral equilibrium. Our central behavioral assumption is that this equilibrium strategy is an ideal free distribution determined by the payoff given by eq. (3). Thus, individuals are able to distribute themselves “freely” over the different foraging distances, such that their payoff from each distance where foraging occurs is the same and at least as high as at distances not visited (Fretwell and Lucas, 1970; Křivan et al., 2008). In a population consisting of *N* birds, the number of individuals foraging at distance *z* from the island is given by *p*(*z*)*N*. Hence, *p*(*z*) can also be interpreted as the proportion of individuals foraging at distance *z*. As birds are assumed to fly with equal probability in all directions, the density of birds foraging at a specific point at that distance equals the proportion of birds foraging at distance *z*, multiplied with the bird’s population size, and divided by the circumference of the circle with radius *z, p*(*z*)*N*/(2*πz*). The rate of prey depletion at that point is the product of this density and the functional response.

Finally, we assume that the average payoff

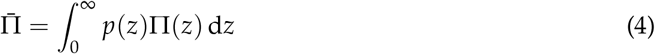

to an individual at the ideal free distribution determines the number of successfully raised offspring. The average payoff thus affects the birds’ population size and hence the number of competitors at any given distance. Without loss of generality, we assume that this payoff is converted into offspring with an efficiency equal to one.

### 2.2 Equilibrium conditions for prey, bird behavior and demography

According to our assumptions, the equilibrium prey density *R*(*z*), the ideal free distribution *p*(*z*), and the equilibrium abundance *N* of birds are determined by the following coupled system of equations.

First, prey density at distance *z* reaches an equilibrium when its renewal is balanced by predation due to foraging birds at that distance. This is the case when

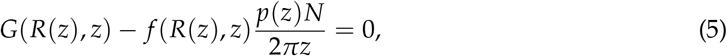

where the second term on the left-hand side is the rate of prey removal at distance *z*, and

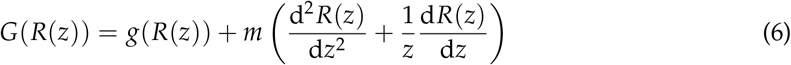

is the prey renewal rate at that distance. The first term on the right-hand side of eq. (6) describes the prey’s growth rate. The second term in eq. (6) describes movement (or “diffusion”), which is proportional to the movement rate *m* and depends on the first and second derivative of the prey density with respect to distance *z* (see Appendix A.2 for why this is so).

Second, the distribution of foraging distances *p*(*z*), as it results from the bird’s behavior, is characterized as an ideal free distribution and thus by the fact that, given individuals in the population follow this behavior, the payoff to all individuals must be identical at all distances where they forage, and must be at least as high as the payoff individuals would obtain at distances where they do not forage. Thus,

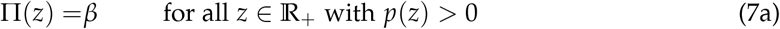

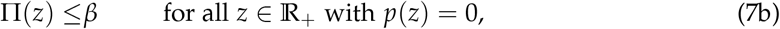

where *β* is the constant payoff obtained from visited foraging distances (the interval of foraging distances satisfying eq. (7a) is the support of the ideal free distribution). Note that, while the payoff given by eq. (7) does not depend explicitly on *p*(*z*) (recall eqs. 1-3), a dependence enters through eq. (5) as the payoff depends on the equilibrium prey density *R*(*z*), which depends on the ideal free distribution expressed by all other individuals in the population. At the ideal free distribution characterized by eq. (7), the payoff obtained from foraging at any visited distance is no less than the payoff obtained from foraging at any possible distance (i.e., Π(*y*) ≤ Π(*z*) for all *y* ∈ *ℝ*_+_ and all *z* ∈ *ℝ*_+_ with *p*(*z*) > 0). This implies that when individuals in the population behave according to the ideal free distribution, no individual has an incentive to unilaterally change behavior and the ideal free distribution is equivalent to a Nash equilibrium (Cressman et al., 2004; Křivan et al., 2008). This makes the ideal free distribution a fundamentally relevant behavioral equilibrium concept.

Third, the forager population is at its demographic equilibrium *N* when the number of successfully raised off-spring is balanced by death, that is, when the average payoff equals the death rate *µ*, which is here assumed to be constant,

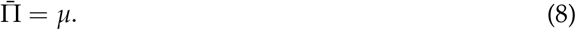

The birth rate as given by the average payoff depends on *N* indirectly through the dependence of the equilibrium prey density *R*(*z*) on *N* (see eq. 5). Note that eq. (7) implies that the average payoff is 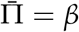 and therefore, due to eq. (8), we have Π(*z*) = *µ* for all *z* with *p*(*z*) > 0.

The coupled system of equations (5)–(8) characterizes the prey equilibrium *R*(*z*), the birds’ behavioral equilibrium *p*(*z*) and their equilibrium abundance *N*. In the following section, we derive explicit expressions for this joint equilibrium.

## 3 Analysis

### 3.1 Single bird species: prey halo and ideal free bird distribution

We start by observing that the flying cost *T*_f_ = 2*z*/*v* monotonically increases with distance *z*. As a consequence, when holding *R*(*z*) fixed, the functional response *f* (*R*(*z*), *z*) (eq. 2), and therefore the payoff Π(*z*) (eq. 3), decrease with *z* and go to zero as *z* becomes large. Hence, a maximal traveling distance *z*_max_ has to exist beyond which prey is left unconsumed, *p*(*z*) = 0 for *z* > *z*_max_. Beyond this maximum, the prey will therefore reach its equilibrium density *R*^*^ in the absence of predation. The payoff at this maximum distance satisfies

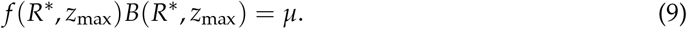

Solving this equation for *z*_max_, using eqs. (1)–(2), gives

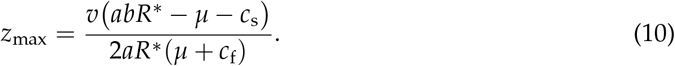

Solving eq. (7) for *R*(*z*), eq. (5) for *p*(*z*) and using the equality *f* (*R*(*z*), *z*) = *µ*/*B*(*R*(*z*), *z*)), the normalization 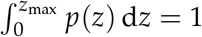 along with Π(*z*) = *µ*, we obtain

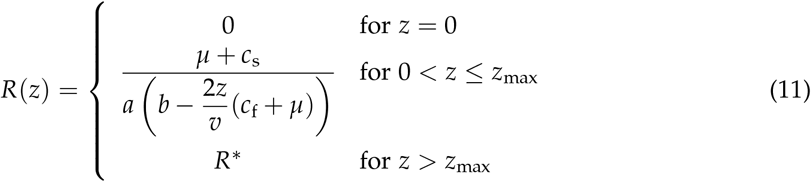

and

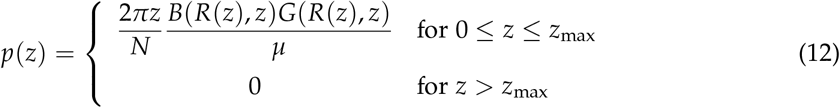

where

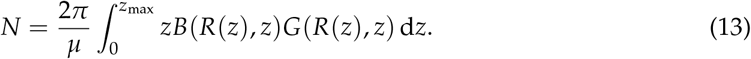

Eqs. (11)–(13) provide an explicit representation of the joint equilibrium for prey, bird behavior and demography, since eq. (11) provides an explicit expression for *R*(*z*), which, in turn, determines *p*(*z*) and *N*. Figure 1(a) illustrates the shape of *R*(*z*) and *p*(*z*) as a function of distance from the island *z*. The fact that we can jointly characterize the equilibrium distribution of prey, bird behavior and demography does not hinge on our specific mechanistic assumptions arising from seabird ecology (i.e., the details on the right-hand sides of eqs. 1, 2, and 6) but holds more generally whenever predators adopt an ideal free distribution along a continuous resource axis. The following conclusions can be drawn. First, eq. (11) shows that the equilibrium prey density *R*(*z*) has its lowest value at *z* = 0, thereafter increases with increasing *z*, and eventually reaches the equilibrium in the absence of consumption, *R*^*^. This is a prey depletion halo, a generic feature of our model. The shape of the halo depends only on the biology of the bird species and thus neither on the details of prey renewal nor movement. This remarkable property has its counterpart in standard predator-prey theory, where the prey equilibrium density only depends on properties of the predator as long as the predator’s functional response is independent of its own density (e.g., Iannelli and Pugliese, 2014, pp. 159). Second, the extent of the prey depletion halo, as given by *z*_max_ (eq. 10), increases with the birds’ flying speed *v*, the prey’s energy content *b*, and the maximum prey density *R*^*^, and decreases with flying and search costs, *c*_s_ and *c*_f_, respectively. Third, the distribution of foraging distances *p*(*z*) starts at *p*(0) = 0, and in the absence of movement (*m* = 0) ends at *p*(*z*_zmax_) = 0 (for an example, see blue curve in fig. 1a).

**Figure 1:**
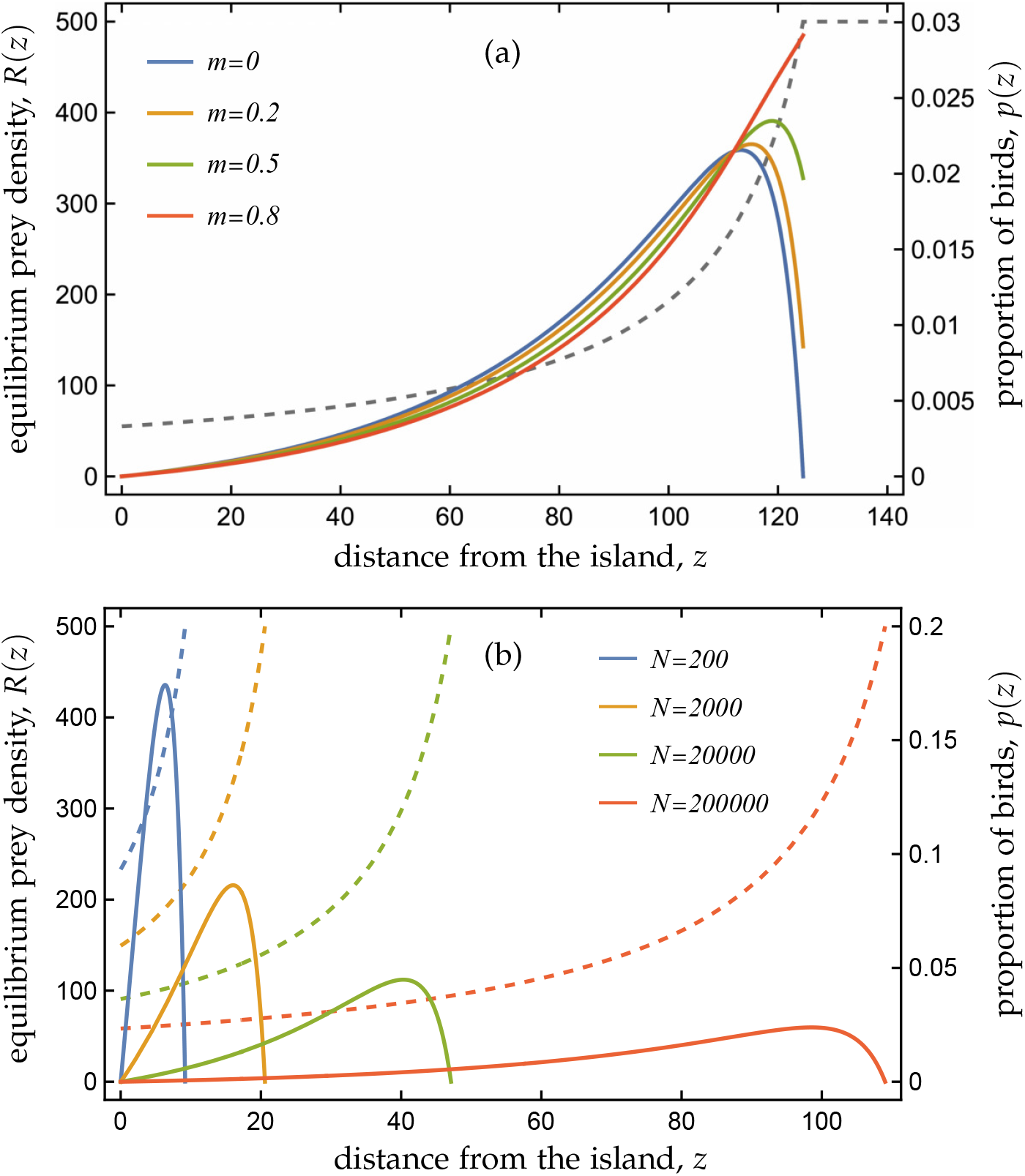
Equilibrium prey density *R*(*z*) (hatched lines, left y-axis) and ideal free distribution *p*(*z*) (proportion of birds feeding at distance *z*, solid lines, right y-axes), as a function of distance *z* from the island under logistic prey growth (*g*(*R*(*z*)) = *R*(*z*)(*r* − *αR*(*z*)), where *r* is the intrinsic per-capita growth rate and *α* the sensitivity to competition), which describes the density-dependent dynamics of a self-renewing prey species. (a) Results under the assumption that birds have reached their demographic equilibrium for four different values of the movement rate *m*. For the parameters given below, in the order of increasing values of the movement rate *m*, the equilibrium colony size *N* equals 288 858, 307 085, 334 427 and 361 768. Note that the shape of the prey depletion halo, given by *R*(*z*), is independent of *m* (cf. eq. 11). At distances *z* > *z*_max_, the prey density is at its equilibrium in the absence of predation, *R*^*^ = *r*/*α* = 500. (b) Results for four different fixed population sizes *N*. Prey depletion halo’s become more pronounced and birds fly further with increasing population size. To avoid overloading the figure, horizontal lines indicating the prey density *R*^*^ at distances *z* > *z*_max_ have been omitted. Parameter values: *r* = 0.02, *α* = 0.00004, *b* = 1, *c*_f_ = 0.2, *c*_s_ = 0.5, *a* = 0.01, *v* = 70, (a) *µ* = 0.05, (b) *m* = 0.

Increasing the prey movement rate increases the value of *p*(*z*_zmax_) (fig. 1a). Hence, higher prey movement results in birds exploiting distances further away from the island more heavily. This is because prey diffusion results in a net movement of prey from distances with higher density to distances with lower density. Fourth, equilibrium colony size *N*, as given by eq. (13), increase with the prey’s net energy content *B*(*R*(*z*), *z*) and its renewal rate *G*(*R*(*z*), *z*). Since *G*(*R*(*z*), *z*) is linearly increasing in the movement rate *m*, so is the birds’ equilibrium population size, as given in the legend of fig. 1(a).

### 3.2 Single bird species at fixed population size

In the above analysis, we assumed that the bird population is at its demographic equilibrium, emphasizing the fact that the area restricted foraging during breeding indeed sets the limit to colony size as hypothesized by Ashmole (1963). However, many seabird populations are not at this equilibrium, for instance, due to human disturbances (e.g., Hughes et al., 2008) or limited nesting sites. In this section, we extend our analysis to the case that the bird population size is fixed for a number *N* that is not equal to the demographic equilibrium. This allows us to make predictions about the extent of prey depletion halos depending on current population size. In this case, the mean payoff *β* has to fulfill the constraint 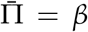 for all *z*, see eq. 7a), where *β* is an unknown quantity. Replacing *µ* with *β* in the expression for the maximum flying distance *z*_max_ given by eq. (10), eq. (4) with *z*_max_ as upper boundary in the integral can be solved for *β*. This results in an implicit equation for *β* that can then be solved numerically given a population size. Once the value of *β* is obtained, eqs. (11) (again, after replacing *µ* with *β*) and (12) provide an explicit solution for the ideal free distribution *p*(*z*) and the equilibrium prey density *R*(*z*). The result is shown in fig. 1(b) for four different population sizes *N*. This figure shows that the maximum flying distance roughly doubles with each ten-fold increase in population size (for the population sizes 200, 2000, 20000, and 200000 the corresponding values of *z*_max_ are 9.2, 20.6, 47.1, and 109.0). This result is in qualitative agreement with the empirically determined distributions shown in fig. 2(b) of Patterson et al. (2022) for two species of murres (*Uria* spp.) when taking from their study the distances up to where 95% of the foraging trips occur.

**Figure 2:**
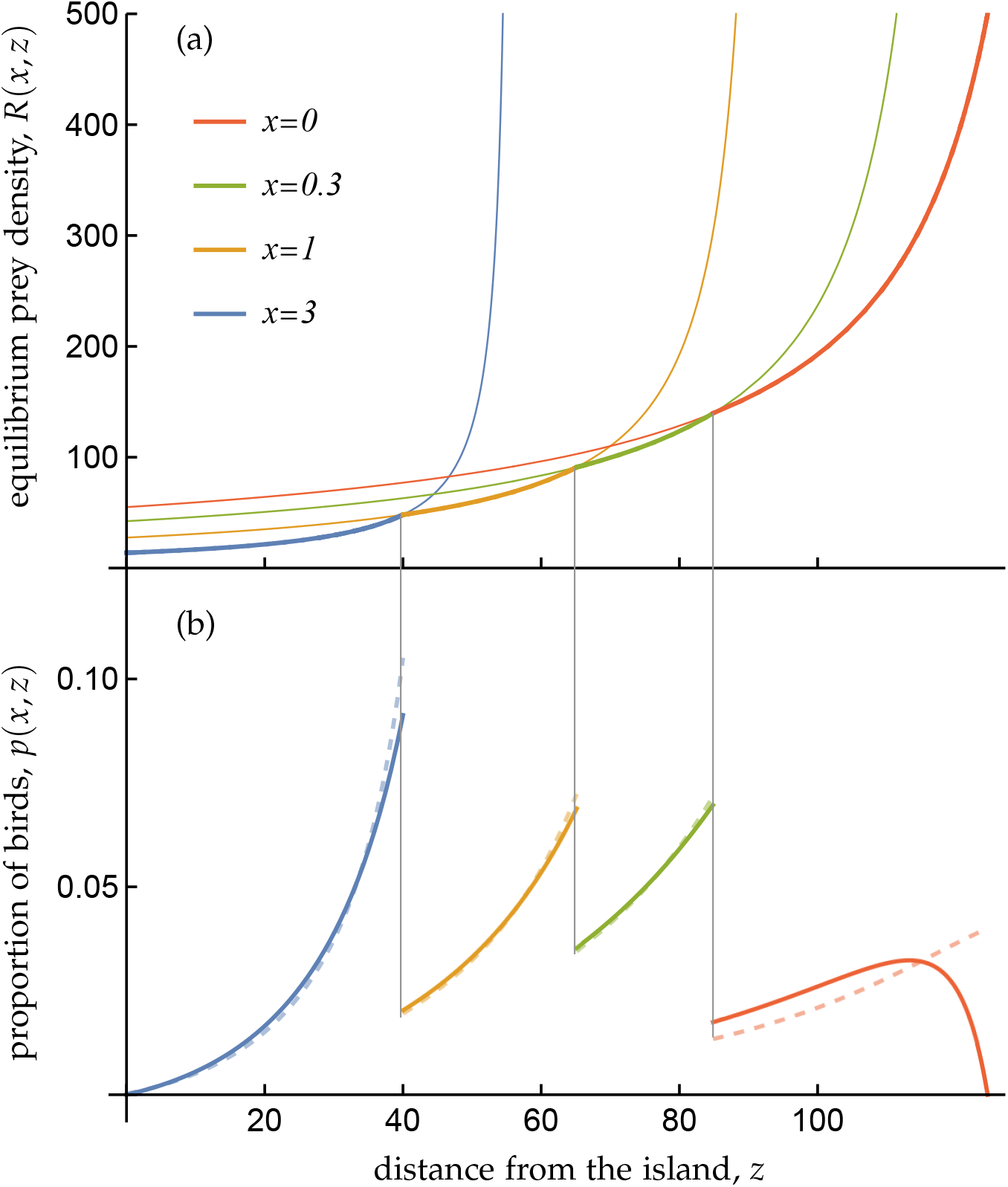
Equilibrium prey density *R*(*x, z*) in (a) and ideal free distribution *p*(*x, z*) (proportion of birds feeding at distance *z*) in (b) as a function of distance *z* from the island for four different trait values *x*. The lowest equilibrium prey density curve is shown as a thick line. Due to competitive exclusion, bird species differing in their trait value *x* only forage at distances where they reduce prey density to a lower value than the other species, resulting in the multi-species ideal free distribution shown in (b) and a multispecies prey depletion halo, given by the composite thick line in (a). Grey vertical lines, connecting panel (a) and (b), are drawn at the *z*-values where species identity changes. The ideal free distribution is shown for two values of the movement rate; solid lines correspond to *m* = 0 and hatched lines to *m* = 0.8. For *m* = 0, the equilibrium population size for the four bird species equal, presented in order of decreasing *x*-values, 8087, 29 531, 51 422, and 192 356. For *m* = 0.8 the populations sizes, following the same order, equal 9941, 32 278, 54 907, and 262 241. Parameter values as in fig. 1(a), except that the values for *a* and *v* are now trait dependent according to the functions *a*(*x*) = *a*_0_ + *a*_1_*x* and *v*(*x*) = *v*_0_/(1 + *v*_1_*x*), respectively, with *a*_0_ = 0.01, *a*_1_ = 0.01, *v*_0_ = 70, *v*_1_ = 0.5.

The payoff *β* at the ideal free distribution for the above population sizes are 1.83, 0.99, 0.41, and 0.08, respectively. This exemplifies Ashmole’s idea (Ashmole, 1963) that the rate of prey delivery to nestlings decreases with increasing population size (as supported by several empirical studies (Hunt et al., 1986; Lewis et al., 2001; Ainley et al., 2004; Ballance et al., 2009; Oppel et al., 2015; Jovani et al., 2016).

We note that for fixed *N* the equilibrium prey density distribution *R*(*z*) is no longer independent of the movement rate *m*. This is illustrated in fig. C1. Higher values of *m* result in a shorter maximal flying distance and a higher prey abundance at each distance *z*. This is in qualitative agreement with simulation results by Weber et al. (2021, their fig. 3B).

**Figure 3:**
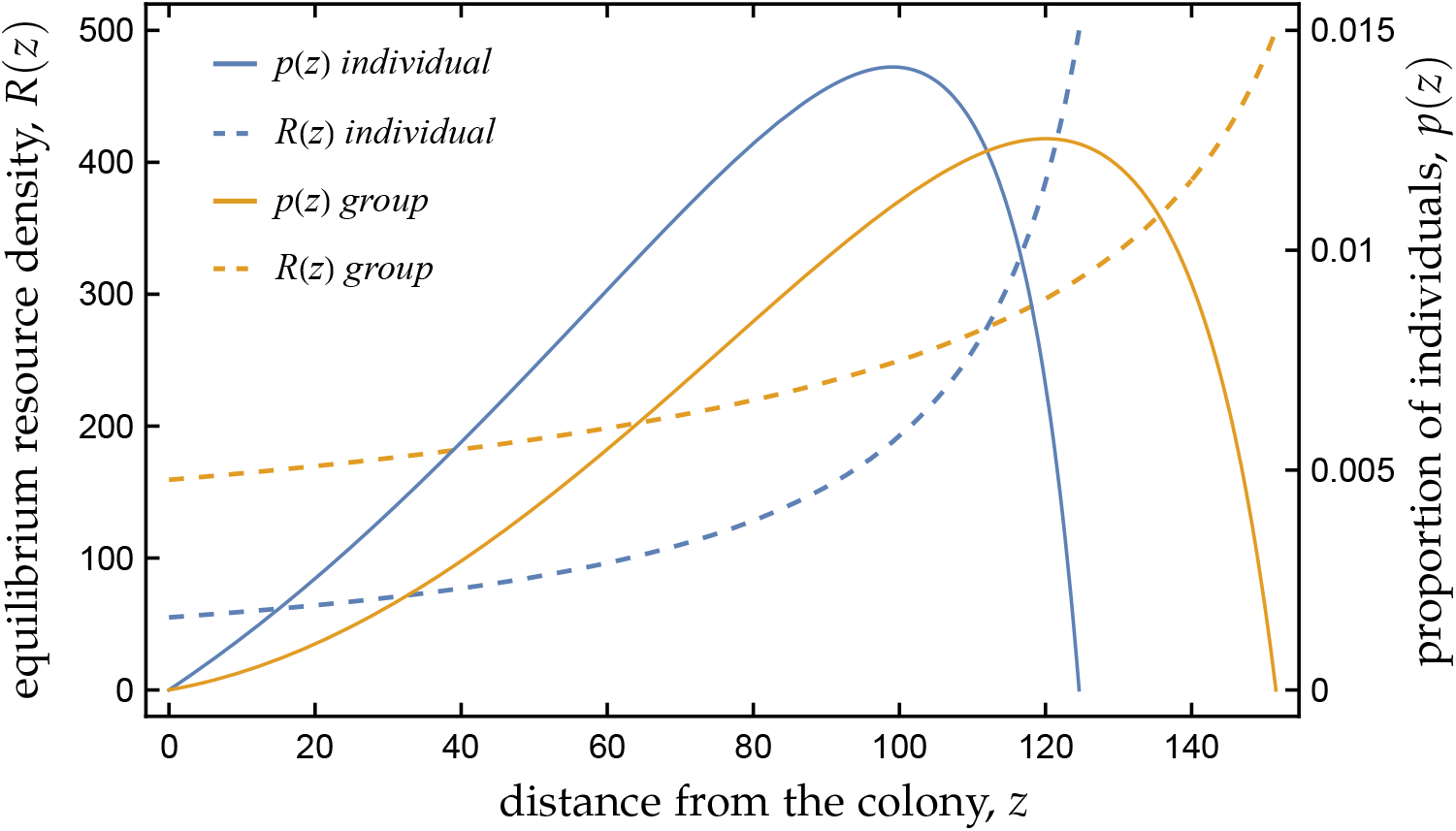
Equilibrium prey density *R*(*z*) (hatched lines, left y-axis) and proportion of individuals foraging *p*(*z*) (solid lines, right y-axes) as a function of distance *z* from the colony for individual (in blue) and group foragers (in orange) under constant resource renewal (“chemostat dynamics”; *g*(*R*(*z*)) = *b*_R_ − *d*_R_*R*(*z*), where *b*_R_ and *d*_R_, are the constant resource renewal and per-capita death rate, respectively). For the used parameters, equilibrium population size of individual foragers equals *N* = 36859, and this value is used as a parameter for group foragers. Hence, given equal population size, group foragers use distances close to the colony less intensively and instead utilize a larger area around the colony (*z*_max_ = 152 for group foragers vs. *z*_max_ = 125 for individual foragers). Note that the two curves for *R*(*z*) and for *p*(*z*) intersect with each other at the same value (*z* = 112). The reason is that the resource equilibrium density at distance *z* is a direct consequence of the number of individuals foraging at that distance. Parameters for chemostat dynamics: *b*_R_ = 0.4 and *d*_R_ = 0.0008. All other parameters as in fig. 1.

### 3.3 Multiple bird species and behavioral niche partitioning

In the above analysis, we considered a single bird species. However, many seabirds breed in mixed colonies, which raises the question of how these species can coexist despite all foraging in the same surrounding waters (Petalas et al., 2024)? A possible answer comes from the following empirical studies. Weber et al. (2021) observed that on Ascension Island, two species of boobies and frigate birds utilize different distances from the breeding colony. Brown boobies (*Sula leucogaster*) forage closer to the colony than its less heavy relative, the Masked booby (*Sula dactylatra*). The Ascension frigatebird (*Fregata aquila*), having a much lower wing-loading (body weight/wing area) than both boobies, forages even further from the colony. Similarly, Razorbills (*Alca torda*) and Common guillemots (*Uria aalge*), breeding in a mixed colony on the Isle of May, Scotland, differ in the distribution of their foraging trips (Thaxter et al., 2010). Razorbills, having a lower wing-loading than Common guillemots, fly on average larger distances. These studies suggest that coexistence may be the result of spatial niche segregation, with some species foraging primarily near the colony while others venture farther away, and that phenotypic differences determine which species occupies each foraging zone.

Here, we formalize this idea by assuming that birds are characterized by a quantitative trait *x* (*x* ∈ ℝ) that can affect their capture efficiency *a*, their flying speed *v*, and their flying and search costs, *c*_f_ and *c*_s_, respectively, to which we refer collectively as foraging components. Different species have different trait values *x* and therefore differ in their foraging components (but we assume that they have the same death rate *µ*), while individuals of the same species share the same trait value *x*. A natural assumption is then that the performance in different tasks is subject to trade-offs. For instance, a morphology allowing for a high flying speed *v* might be less efficient at foraging by plunge-diving and therefore would be coupled to a lower capture efficiency *a*. For the Alcidae (auks), a group of seabirds catching prey by wing-propelled diving, it has been proposed that smaller wings, resulting in a high wing-loading, increase flight costs *c*_f_ but make for better divers (Gaston, 2004; Elliott et al., 2013), resulting in higher capture efficiency *a*. This trade-off has been confirmed by Thaxter et al. (2010), who demonstrate that Razorbills are poorer divers compared to Common guillemots.

In the following, we show that such trade-offs indeed allow for the coexistence of multiple seabird species through a partitioning of the waters surrounding the colony into distinct circular foraging zones. In this analysis, we assume that each bird species reaches its demographic equilibrium (as in Section 3.1).

In order to indicate that the foraging components can differ with *x*, we henceforth write these as functions of *x*, for instance, *a*(*x*). Similarly, we add the argument *x* to the payoff function (eq. 3, Π(*x, z*)), the maximum flying distance (eq. 10, *z*_max_(*x*)), the equilibrium prey abundance (eq. 11, *R*(*x, z*)), the ideal free distribution (eq. 12, *p*(*x, z*)), and the equilibrium bird population size (eq. 13, *N*(*x*)) to indicate that these functions depend on the birds’ phenotype.

To investigate how trait differences allow for coexistence, we start by observing that a species with trait value *x* induces a corresponding equilibrium prey density curve *R*(*x, z*) (in the absence of any other species). Figure 2(a) shows such curves for four trait values *x* under the assumption that capture efficiency and flying speed are negatively correlated (*a*(*x*) is a monotonically increasing function while *v*(*x*) is a monotonically decreasing function in *x*). This figure shows that for any two species, the curves *R*(*x, z*) intersect exactly once. For illustration, let us focus on two species, say, those characterized by *x*_1_ = 3 (henceforth called species 1, blue curve) and *x*_2_ = 1 (henceforth called species 2, orange curve). Upon examination of the equilibrium prey density curves induced by species 1 and 2, denoted *R*(*x*_1_, *z*) and *R*(*x*_2_, *z*), respectively, we can draw the following conclusions. First, in the interval between the island (*z* = 0) and the point where the two equilibrium prey density curves intersect, species 1 depletes prey to a lower density compared to species 2. We denote the distance to this intersection point with *z*_1_, which in our example is approximately 40. Second, in the interval between *z*_1_ ≈ 40 and the maximum flying distance of species 2, *z*_max_(*x*_2_), the situation reverses. Here, species 2 depletes prey to a lower density than species 1. In short, *R*(*x*_1_, *z*) < *R*(*x*_2_, *z*) for *z* ∈ [0, *z*_1_] and *R*(*x*_1_, *z*) > *R*(*x*_2_, *z*) for *z* ∈ [*z*_1_, *z*_max_(*x*_2_)]. The fact that *R*(*x*_1_, *z*) < *R*(*x*_2_, *z*) in the interval [0, *z*_1_] implies that the payoff from foraging at these distances to species 2 is lower than the death rate (Π(*x*_2_, *z*) < *µ*). Then, according to eq. (7b), species 2 does not utilize this interval (*p*(*x*_2_, *z*) = 0 for *z* ∈ [0, *z*_1_]). The reverse is true for the interval [*z*_1_, *z*_max_(*x*_2_)]. Here, Π(*x*_1_, *z*) < *µ* so that species 1 does not utilize this interval (*p*(*x*_1_, *z*) = 0). In conclusion, we find a partitioning of the surrounding of the colony where species 1 only exploits distances close to the island (*z* ∈ [0, *z*_1_]) and species 2 only exploits distances further away from the island (*z* ∈ [*z*_1_, *z*_max_(*x*_2_)]). In other words, the species with the trait value inducing the lowest equilibrium prey density at a certain distance *z* excludes species with other trait values from foraging at that distance. Our preceding argument shows that for two species to be able to coexist, it is a necessary (and sufficient) requirement that the corresponding equilibrium prey density curves *R*(*x, z*) intersect with each other. This argument, which is developed more formally in Appendix C.3, echoes the resource-ratio rule of community ecology. It states that if multiple species compete for a single limiting resource, here the resource at distance *z*, then whichever species can survive at the lowest equilibrium resource level outcompetes all others (Tilman, 1982; Grovers, 1997), resulting in competitive exclusion (Levin, 1970).

More generally, in Appendix C.2, we show that any two equilibrium prey density curves *R*(*x, z*) intersect at most once. From this follows, that two curves *R*(*x, z*) intersect with each other if and only if the values of the equilibrium prey density at distance *z* = 0 (*R*(*x*, 0)) and the maximum flying distance *z*_max_(*x*) have the same ordering. For our example above, fig. 2 shows that *R*(*x*_1_, 0) < *R*(*x*_2_, 0) and *z*_max_(*x*_1_) < *z*_max_(*x*_2_). Thus, the trade-off between foraging components induces a trade-off in the efficiency to forage at short and long distances from the colony. Species 1, having a higher capture efficiency, is more efficient at exploiting prey close to the colony, resulting in a lower equilibrium prey density at *z* = 0, at the cost of having a shorter maximum traveling distance, while for species 2, having a higher flying speed, the situation is reversed.

In Appendix C.3, we show that the above arguments generalize to any finite community *χ* = {*x*_1_, *x*_2_, …, *x*_*n*_} with *n* > 2 bird species (fig. 2 shows it for *n* = 4). Thus, whenever we have coexistence of a set of species, they forage in mutually exclusive distance intervals around the island (until a distance is reached beyond which no individual forages), each species occupying exactly one interval. In each interval, the prey distribution is determined by the foraging components, and thus the trait value *x*, of the species occupying that interval according to eq. (11). The corresponding ideal free distribution and abundance of that species are fully characterized analytically and given by analogues of eqs. (12)–(13), where the ideal free distributions of each species ranges over an interval that depends on the trait values of all other species in the community (see eqs. C14–C16). The result is a multi-species prey depletion halo, as shown in fig. 2(a), and a multi-species ideal free distribution, as shown in fig. 2(b). Hence, our main finding is that, in the presence of trade-offs between foraging components, multiple bird species can coexist on a single prey species by partitioning the area around the colony into discrete distance intervals, a form of behavioral niche portioning. Just as in the single-species case, prey movement does not affect the equilibrium prey distribution. As a consequence, prey movement does not affect the partitioning of distances of coexisting bird species. However, just as in the single species case, prey movement results in a shift of the ideal free distribution of each species towards higher distances due to a net movement of prey individuals toward distances with lower prey density (see fig. 2b).

Figure 2 is based on the assumption that capture efficiency *a*(*x*) and flying speed *v*(*x*) are negatively correlated. In Appendix D, we determine for all possible pairwise trade-offs between foraging components whether they have the potential to induce a trade-off between *R*(0, *x*) and *z*_max_(*x*), which is the prerequisite for species coexistence (allowing for more than two foraging components to vary with *x* does not result in qualitative new results). The result is summarized in table 1. Two different outcomes occur. First, a trade-off between foraging components does not induce a trade-off in the efficiency to forage at different distances. This case is illustrated in fig. D1. Second, whether a trade-off between foraging components induces a trade-off between *R*(0, *x*) and *z*_max_(*x*) depends on parameter and trait values. Figure 2 shows an example where indeed four species can coexist, while fig. C2 shows an example where coexistence is possible for some species but not for others.

**Table 1:** The four foraging components capture efficiency *a*, flying speed *v*, flight costs *c*_f_ and search costs *c*_s_ allow for six different pairwise combinations. For each combination, we determine whether a negative correlation between the foraging components has the potential to mediate coexistence of species with different trait values *x*. See Appendix D for details.

| coexistence | $a \ \& \ v$ | $a \ \& \ c_f$ | $a \ \& \ c_s$ | $v \ \& \ c_f$ | $v \ \& \ c_s$ | $c_f \ \& \ c_s$ |
| --- | --- | --- | --- | --- | --- | --- |
| parameter-dependent | ✓ | ✓ |  |  | ✓ | ✓ |
| not possible |  |  | ✓ | ✓ |  |  |

The results from Table 1 can be understood as follows. Foraging comes with two costs (see eq. 1), namely costs associated with flying (*c*_f_2*z*/*v*) and costs associated with searching for prey (*c*_s_/(*aR*(*z*)). Flight costs increase and search costs decrease with distance *z* from the island. The latter is a consequence of the prey depletion halo around the island. Different species can coexist if they are specialized to forage on different distances from the island by either experiencing low costs when foraging on depleted prey close to the island (high *a* or low *c*_s_) at the cost of higher costs when foraging on less depleted prey further away from the island (low *v* or high *c*_f_) or vice versa (high *v* or low *c*_f_ at the cost of low *a* or high *c*_s_).

## 4 Discussion

Inspired by the evidence provided by Weber et al. (2021) for (i) a prey depletion halo around the seabird colony on Ascension Island and (ii) the spatial partitioning of the feeding grounds surrounding the colony between coexisting seabirds, we here present a comprehensive mathematical model studying the ideal free distribution of foraging trips of central place foragers exploiting a single prey species in a two-dimensional environment. By analytically solving the joint equilibrium of the prey distribution, the behavior of the forager and its demography, we show that a prey depletion halo is a robust outcome of central place foraging. Its shape, given by eq. (11) and illustrated in figs. 1 and 2, is robust in the sense that it is independent of the specifics of the prey population dynamics, including its movement rate, as all parameters in eq. (11) describe properties of the forager.

Our full model includes an arbitrary number of species, where, due to species-specific morphological and physiological trait values, individuals can differ in flying speed, capture efficiency, travel and search costs. We show that coexistence is a robust outcome that emerges whenever species differences result in some species experiencing lower costs when foraging on a prey at low density close to the breeding colony, while others experience lower costs when flying long distances to forage on a prey at high density. A qualitatively similar result has been found by Bolin et al. (2018) in a model describing the foraging behavior of solitary bees, assuming that species vary along a trade-off of low flight costs and efficient resource use. We want to stress that, in contrast to Bolin et al. (2018), coexistence in our model occurs in the presence of a single biological prey species. As explained above, the key to understand this coexistence is that for different species the same prey — but at different distances from the colony — comes with different costs. The result is a multi-species ideal free distribution (fig. 2b) and a multi-species prey depletion halo (fig. 2a), where no two bird species forage at the same distance. Instead, different species exploit mutually exclusive circular zones around the island.

The sorting of species into mutually exclusive foraging zones occurs despite the fact that, in principle, the area close to the colony would be preferred by all species. It is in this vicinity that catching prey comes at the lowest travel costs, both in terms of the time needed to reach the foraging area and the energy investment to do so. Thus, a community of coexisting species that are central place foragers, all feeding on the same prey species, is an example of a community with shared preferences. This is significant since theory in community ecology is dominated by models with distinct preferences, despite indications that shared preference might be the more common scenario (Rosenzweig, 1991; Wisheu, 1998). With distinct preferences, each species performs best in a different part of the niche space, regardless in which community they occur. In our model, where all species perform best close to the island in the absence of competing species, it is in the presence of competitors that deplete prey to such low values that the immediate surrounding of the colony is not sufficiently profitable to be used by species that are less efficient foragers. Thus, for each species, the fundamental niche consists of the circular area around the colony that is limited by the species specific maximum flying distance. However, in the presence of competitors, the realized niche of each species consists of the circular zone around the island where that species is able to deplete the prey to a lower density than any other species.

A consequence of the fact that coexistence results from behavioral niche partitioning is that the conditions for coexistence are relatively mild compared to models where coexistence is only mediated by trait differences without linked behavioral differences. A corresponding result has been found in a model where consumers compete for two resources and where individuals can choose to either attack or ignore a prey item, depending on their expected energy gain from doing so, which depends both on their phenotype and on the abundance of resources as determined by the competitors (Rueffler et al., 2007). In that model, optimal diet choice greatly enlarges the set of pairs of trait values that allow for coexistence. Furthermore, coexistence in our model is generally not a transient phenomenon if one allows for evolutionary change in the trait values *x* that characterize the different species. This can be seen from fig. 2. Changes in the trait value *x* of the four coexisting species changes the pattern of intersections of the equilibrium prey density curves only quantitatively but not qualitatively. Thus, coexistence does not easily break down under evolutionary change. In fact, the opposite is likely to be true. An input of individuals with different trait values, either by mutation or immigration, can readily increase diversity (see fig. C2 for an example).

Weber et al. (2021) report the distribution of seabirds both as the proportion of time birds spend foraging at different distances from the island (their fig. 2A) and as density (their fig. 2B). The first corresponds to the ideal free distribution, as shown in fig. 2(b). Obviously, their distribution markedly differs from ours. For several reasons, this is not surprising. First, our derivation assumes that foragers are “ideal”, that is, they have complete information about both their environment and their own abilities. This is certainly not true for natural systems, where individuals have to build up this information through constant probing (as documented in the Northern gannet (*Morus bassanus*), Votier et al., 2017). Second, intraspecific phenotypic variation occurring in natural populations, both within (Sommerfeld et al., 2013) and between the sexes (Weimerskirch et al., 2009), causes that different individuals of the same species differ in their foraging decisions, while in our model all individuals of a species are identical. Even for a monomorphic species it has been documented that the sexes can differ in their foraging decisions (Lewis et al., 2022). Third, Weber et al. (2021, Table S1) show that, for their investigated seabird species, small differences in diet composition do occur, and Ascension frigatebirds to some extent forage by kleptoparasitism, both complications not present in our model. Fourth, Weber et al. (2021) suggest that the prey depletion halo around Ascension Island expands during the breeding season. Hence, their system is not at equilibrium, while our analysis indeed assumes that the system has equilibrated. In Appendix B, we present results showing how the ideal free distribution of foraging birds translates into bird densities at different distances. We consider three different functions for the prey growth rate, including one where we extend our model to the case that the prey’s growth rate depends directly on distance from the island, describing a gradient due to distance-dependent primary productivity. We show that the shape of the distribution of bird densities is both quantitatively and qualitatively sensitive to the specifics of the growth function, and empirical predictions thus seem only possible given knowledge of empirical details.

It is noteworthy that our model is agnostic as to whether the ideal free distribution is composed of individuals that all distribute their foraging trips according to the ideal free distribution (mixed strategy) or of individuals that each utilize a single distance and that the frequency distribution of individuals utilizing the different distances follows the ideal free distribution (mixture of pure strategies). This pure-mixed strategy equivalence occurs since what determines energy intake at a given distance is the realized density of individuals at that distance, and that at the ideal free distribution the net energy gain is the same at all visited distances. In a review, Ceia and Ramos (2015) report that in seabirds, individual specialization in foraging strategies has been documented in 87% of the studies investigating this phenomenon, suggesting that the ideal free distribution is more likely the result of a mixture of pure strategies. In the absence of diffusion, our model should also be agnostic as to whether prey individuals move in response to predation. This is because the equilibrium prey density *R*(*z*) can be seen as the product of the total prey population size times the proportion that reproduces at distance *z*. This proportion, in turn, can be interpreted as an ideal free distribution for prey reproducing at different distances from the island when payoff at distance *z* is taken to be the per-capita growth rate at that distance that equalizes at all distances. Hence, we suggest that our results can also be read as characterizing the joint predator-prey behavioral and demographic equilibrium.

Finally, we want to emphasize that our results are obtained under the assumption that birds act as individual payoff maximizers. Central place foragers that are selected to maximize group payoff, such as colonies of social insects consisting of highly related individuals, do not follow an ideal free distribution (see fig. 4). Instead, we show that colony payoff is maximized by increasing maximum travel distance and simultaneously underexploiting the vicinity of the colony.

**Figure 4:**
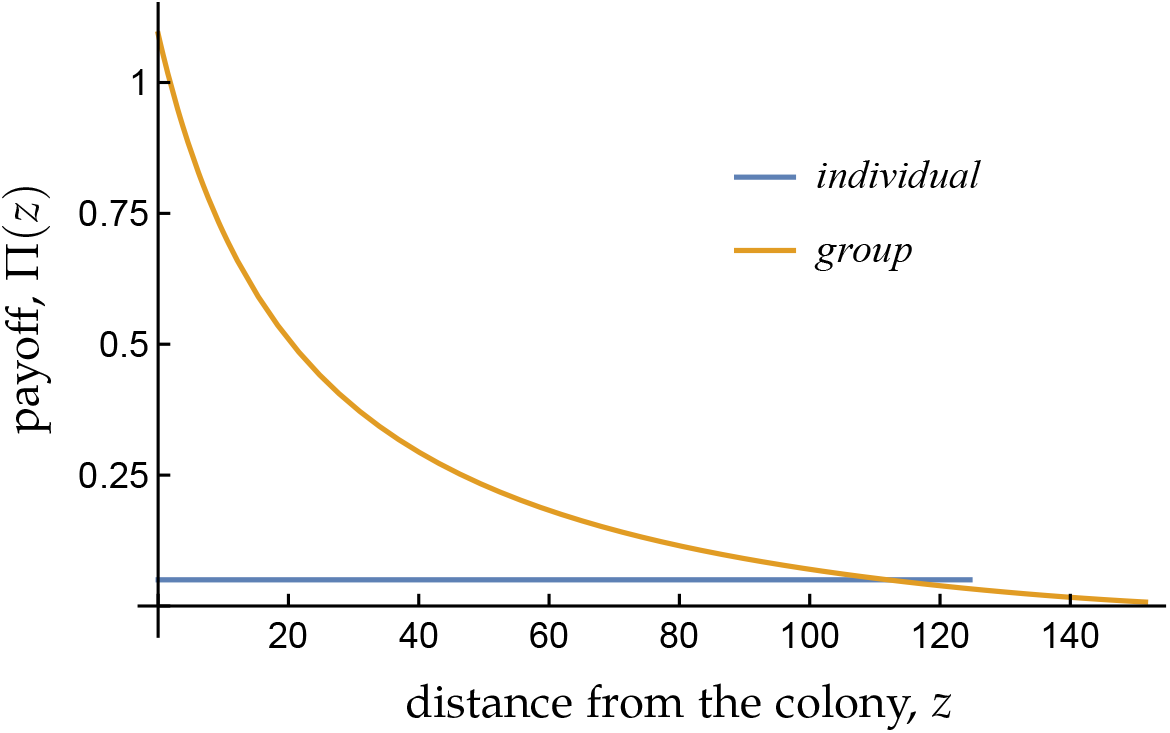
Payoff Π(*z*) for group (in orange) and individual (in blue) foragers. As a result of the ideal free distribution, individual foragers have an identical payoff at all distances (equal to *µ* = 0.05, up to *z*_max_ = 125). In contrast, group foragers under-exploit resources at short distances and over-exploit resources at large distances (fig. 3). As a result, individual payoff is monotonically decreasing with distance *z* (up to *z*_max_ = 152). The two payoff curves Π(*z*) intersect with each other at the same distance as the equilibrium resource curves (*z* = 112, cf. fig.3), as payoff is a direct consequence of resource abundance. Parameters as in fig. 3.

### 4.1 Conclusion

In his influential study of seabirds on Ascension Island, Ashmole (1963) proposed that seabirds breeding in large colonies locally reduce their prey, generating a prey depletion halo around the colony. Strong evidence for such halos has been provided by Weber et al. (2021), a study that further suggests that coexistence of seabirds is made possible by individual foraging decisions resulting in a spatial segregation into different circular zones around the colony. Our model, which is based on features shared by many seabirds breeding in large colonies without incorporating the details of any specific colony, shows that prey depletion halos are a robust outcome, suggesting they should generally be associated with central place foraging and extend beyond seabird ecology. We also show that trait mediated behavioral niche partitioning results in robust coexistence. We suggest that this process facilitates species coexistence compared to standard theory based on trait-mediated niche partitioning as developed in the wake of MacArthur and Levins (1967)’s seminal work. Our trait-dependent multi-species ideal free distribution–a Nash equilibrium of a population game–is not limited to central place foragers, and holds for other predator-prey systems whenever predators adopt an ideal free distribution along a continuous resource axis and the functional response does not depend on predator density. Our results may thus play a more general role in ecology as a coexistence promoting mechanism.

#### Box 1

Group versus individual forages

The results in the main section are derived under the assumption that selection acts to maximize individual payoff. For highly related social insects, however, it is appropriate to assume that there is no conflict between group members and selection acts to maximize colony payoff. Here, we consider a central-place foraging model for such foraging groups, where we search for the behavioral strategy *p*(*z*) that maximizes the net per-capita rate of energy delivery of group members. Our results are based on a non-self-renewing resources, such as nectar in flowers. The equilibrium behavior *p*(*z*) maximizing group payoff is derived in Appendix E, using all biological assumptions of the individual foraging model, except that (i) group size is *N* fixed, and (ii) movement of resources is absent (*m* = 0). distance from the colony, *z*

We find that at the group foraging equilibrium, the maximum travel distance is larger and the distribution of travel distances is more skewed toward large distances compared to the equilibrium for individual foragers (fig. 3). Thus, group foragers exploit regions close to the colony less intensely and instead use a larger foraging area. As a result, group foragers do not equalize payoff from different distances but distribute themselves such that payoffs at short distances exceeds payoffs at long distances (fig. 4), generalizing results from two-patch models by Brown (1998), Dukas and Edelstein-Keshet (1998), and Morris et al. (2001), and from a simulation study by Robinson et al. (2022). This can be explained by the avoidance of kin competition at distances close to the colony where travel costs are lowest. By exploiting resources at large distances, group foragers decrease competition for relatives foraging close to the colony. The increase in payoff due to this decrease in competition more than outweighs the decrease in payoff suffered by individuals from increased competition far away from the colony and by individuals flying further than the maximum travel distance of individual foragers. This results in higher average payoff under group foraging (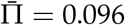compared to 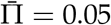 under individual foraging). Thus, group foragers exploit resources more efficiently. Both these trends, decreases in payoff with distance and higher average payoff, are expected to be robust outcomes under group foraging, since they are both direct consequences of the reduced competition entailed by maximizing group payoff.

## Appendix A Model components

Throughout this appendix, our notation makes explicit that each bird is characterized by a quantitative trait *x* ∈ ℝ and is part of a community consisting of *n* different species whose trait values are collected in the set *χ* = {*x*_1_, *x*_2_, …, *x*_*n*_}.

### Appendix A.1 Rate of energy delivery by birds

We here derive the payoff function given by eq. (2) in the main text. Payoff is defined as the net rate of energy delivery to its nestlings of an individual foraging at distance *z*. A foraging trip consists of two activities, flying from the island to a foraging area at distance *z* and back, and searching for prey at that distance. We here assume that birds fly back to the island after catching *n*_L_ prey items, that is, they are *n*_L_-prey loaders. All results in the main text are for single-prey loaders (*n*_L_ = 1). We here show that this does not restrict the generality of the model. The time it takes an individual with trait value *x* to fly from the island to distance *z* and back is *T*_f_(*x, z*) = 2*z*/*v*(*x*) and *T*_s_(*x, z*) = *n*_L_/(*a*(*x*)*R*(*z*)) is the average time for that individual to successfully catch *n*_L_ prey items when foraging at distance *z*. This follows from the rule for compounding exponential distributions, since the mean waiting time for a single catch is exponentially distributed with mean 1/(*a*(*x*)*R*(*z*)).

The total time per foraging trip is thus *T*_f_(*x, z*) + *T*_s_(*x, z*) and let *E*(*x, z*) be the net amount of energy gained during such a foraging trip. The net energy gain per time unit is then

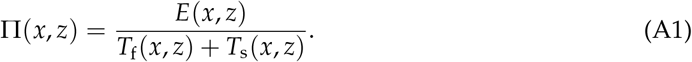

According to our assumptions (section 2.1), birds catch prey at rate *a*(*x*)*R*(*z*), where *a*(*x*) is the capture efficiency of an individual with trait value *x* and *R*(*z*) the equilibrium prey density at distance *z*. Furthermore, each prey item contains on average *b* energy units. The net amount of energy gained during a foraging trip is then given by

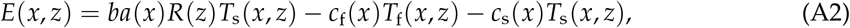

where *c*_f_(*x*) and *c*_s_(*x*) are the energy costs per time unit spend flying and foraging, respectively.

Substituting the expression for *E*(*x, z*), *T*_f_(*x, z*) and *T*_s_(*x, z*) into eq. (A1), we obtain

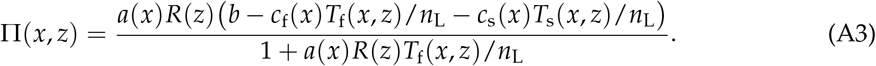

This can be rewritten as

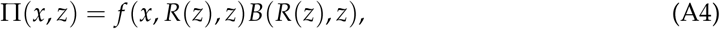

where

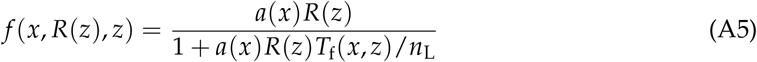

is the functional response of an individual foraging at distance *z*, and

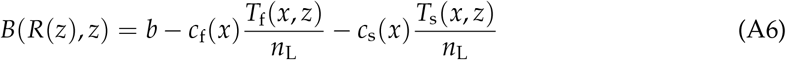

is the net energy content of a prey item from that distance. For *n*_L_ = 1 and dropping the explicit dependence on the trait *x* in all functions results in eq. (2) in the main text. Note that *n*_L_ > 1 does not qualitatively affect the functional form of the model, since it scales the flying and search times equally.

### Appendix A.2 Reaction-diffusion process for the prey

We here derive eq. (5). We assume that the prey follows a radially symmetric reaction-diffusion process with homogeneous diffusion, and ignore reflecting boundary effects of the island, which is assumed to be a single point in space where the prey density is assumed to be zero. We then assume that the partial differential equation describing the dynamics of the density of prey *R*(*t, z*) at time *t* is defined for all *z* ∈ (0, ∞) according to

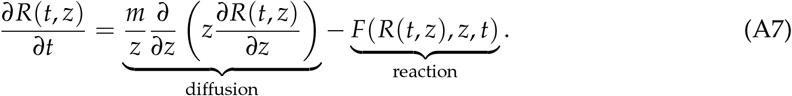

The first term on the right-hand side represents diffusion, occurring at rate *m*, to which we refer in the main text as movement rate. The second term represents reaction, which is some function of prey density, possibly distance from the island and time. Eq. (A7) corresponds to the twodimensional reaction-diffusion process given by eq. (40) in Edelstein-Keshet (1988, Chapter 9).

But since the process is radially symmetric, it is written here in polar coordinates using eq. (61) in Edelstein-Keshet (1988, Chapter 9). The reaction term *F*(*R*(*t, z*), *z, t*) is detailed below. Note that for a reaction term *F*(*z*) that is independent of prey density and time, representing fixed consumption, eq. (A7) is equivalent to the consumption-dispersion model of Weber et al. (2021, their eq. 5).

At equilibrium, *R*(*t, z*) = *R*(*z*) and thus *∂R*(*t, z*)/*∂t* = 0 and *∂*^*n*^*R*(*t, z*)/*∂z*^*n*^ = d^*n*^ *R*(*z*)/ d*z*^*n*^.

Next, let us assume that the reaction term at such an equilibrium is given by

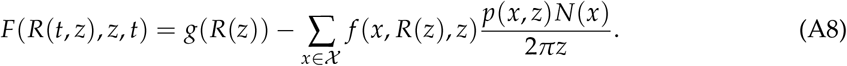

The right-hand side is time-independent and the sum of a density-dependent prey growth rate *g*(*R*(*z*)) at distance *z* and the depletion rate *f* (*x, R*(*z*), *z*)*p*(*x, z*)*N*(*x*)/(2*πz*) due to birds foraging at distance *z*, summed over all bird species with trait values in *χ* = {*x*_1_, *x*_2_, …, *x*_*n*_}. Here, *p*(*x, z*)*N*(*x*)/(2*πz*) is the density of individuals with trait value *x* at distance *z*. The denominator 2*πz* accounts for the fact that individuals foraging at distance *z* distribute themselves randomly over the circumferences of the circle centered at the island and *N*(*x*) is the number of individuals for the species with trait value *x*. Hence, at equilibrium, eq. (A7) can be written as

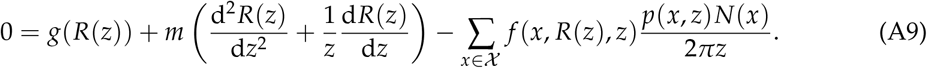

Let us define

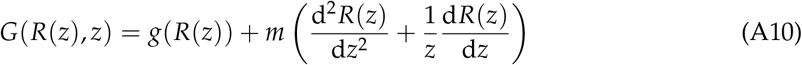

as the endogenous rate of change of prey density at distance *z*, called renewal rate in the main text. Assuming that the bird community consists of only a single species (*χ* = {*x*}), so that the argument *x* can be dropped in all relevant functions, and substituting *G*(*R*(*z, z*) into eq. (A9), we obtain eq. (5) in the main text.

In the absence of consumption, the last term on the right-hand side of eq. (A9) equals zero. We assume that the resulting equilibrium prey density is homogeneous over space and denote it as *R*^*^. This satisfies eq. (A9) with *g*(*R*^*^) = 0.

## Appendix B The single species ideal free distribution expressed as density

In their fig. 2B, Weber et al. (2021) present the empirically determined distributions of foraging birds around Ascension Island – when expressed as densities. In this appendix, we present results from our model about how the density of birds (number of birds/area) foraging at distance *z* from the island changes with distance. We do this for the case of a single bird species and thus drop the argument *x* from all quantities.

The density of birds at distance *z* is given by

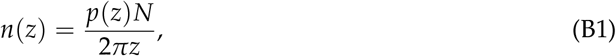

which, from eqs. (12)–(13), can be written as

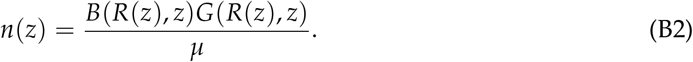

The shape of this function depends on the shape of the prey renewal rate *G*(*R*(*z*), *z*), which is equal to the prey growth rate *g*(*R*(*z*)) in the absence of prey movement (*m* = 0, eq. 6). To illustrate how different biological assumptions about the prey’s growth rate affect the density of birds feeding at a given distance, we consider three different growth functions.

### Logistic growth

Logistic growth describes the density-dependent dynamics of a self-renewing prey and can be written as

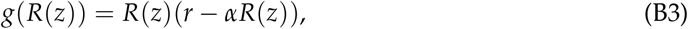

where *r* is the intrinsic per-capita growth rate and *α* the sensitivity to competition. The righthand side of eq. (B3) is a quadratic function in *R*(*z*), representing a parabola that is open to the bottom, intersects with the x-axis at *R*(*z*) = 0 and *R*(*z*) = *r*/*α*. The growth rate of the prey population increases with population density as more individuals can produce more offspring, but this is counteracted by increased negative density dependence. These two opposing forces cause that the growth rate is maximal when the prey density equals half the equilibrium density, *R*(*z*) = *r*/(2*α*). We discuss the effect of logistic prey dynamics on the density of birds after we introduced the next growth function.

**Figure B1:**
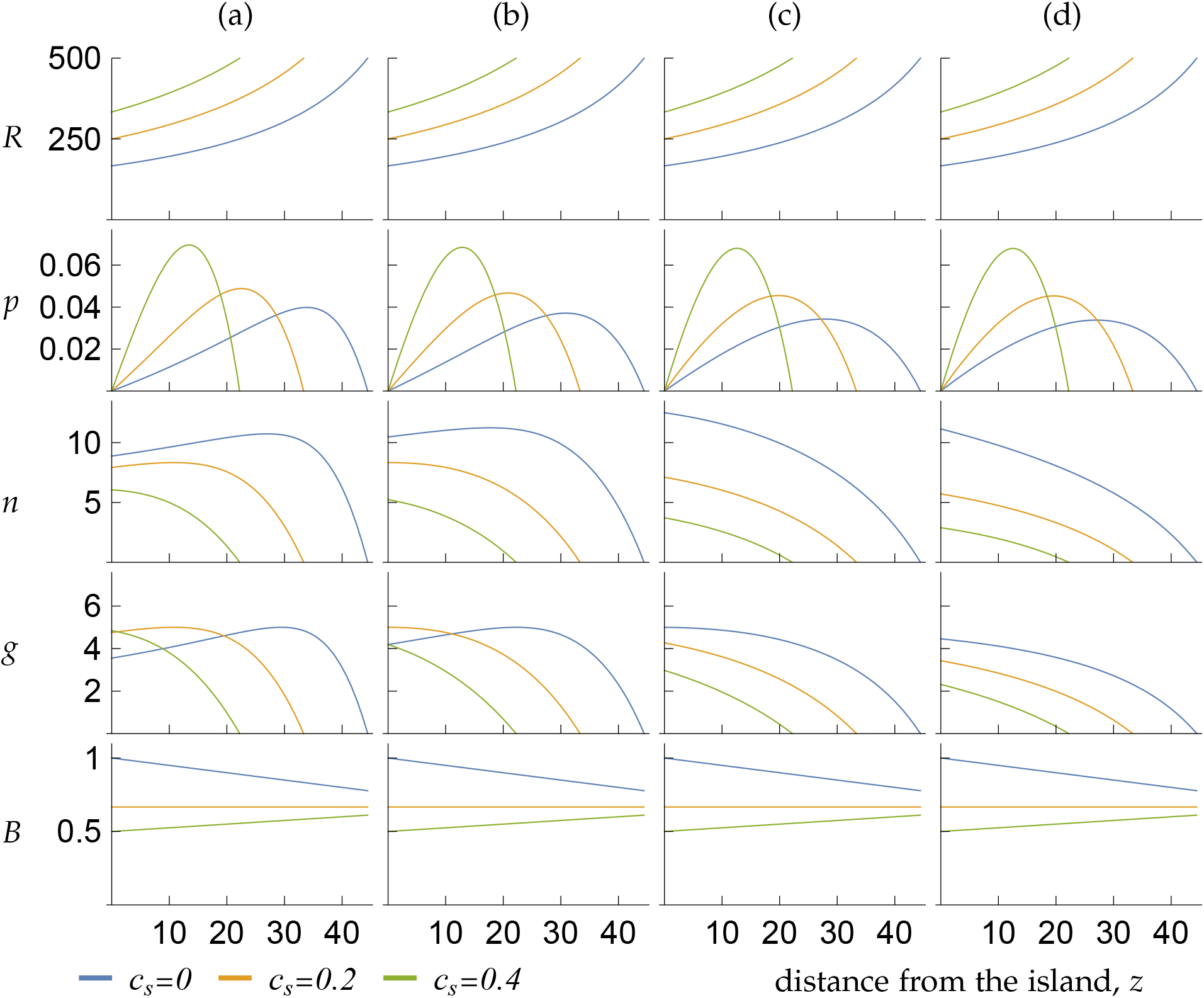
Equilibrium prey distribution *R*(*z*) (first row), proportion of birds *p*(*z*) (second row), bird density *n*(*z*) (third row), prey renewal rate *G*(*R*(*z*), *z*) (fourth row), and net energy content *B*(*R*(*z*), *z*) (fifth row) as a function of distance *z* from the island under the maturation model for the four different growth functions *g*(*R*) shown in fig. B2 (in columns). Note that the panels within the first and last row are identical to each other, since neither the equilibrium prey density nor the net energy content depend on *g*(*R*(*z*)) (eqs. 11 and 1, respectively). Parameter values: *m* = 0, *b* = 1, *c*_f_ = 0.2, *a* = 0.0024, *v* = 80, *µ* = 0.4.

**Figure B2:**
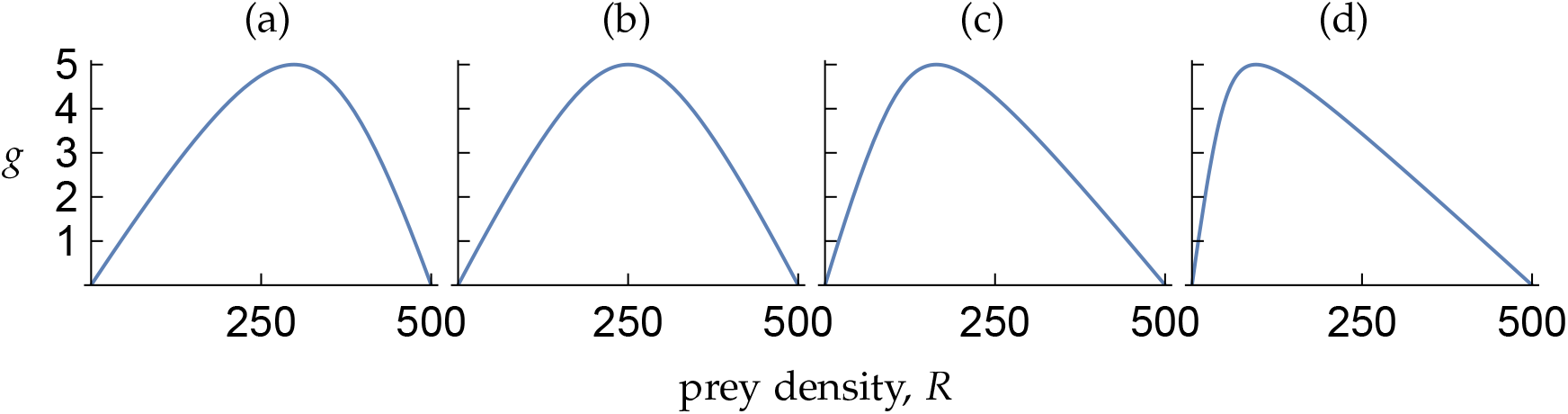
Growth rate *g*(*R*) of adult prey as a function of adult density *R* for four parameter combinations of the maturation model (eq. B8). Parameters are chosen such that the equilibrium density of adult prey in the absence of predation by birds equals 500 and the maximum growth rate equals 5. (a) *b*_*R*_ = 1.5*d*_*R*_: The maximum growth rate occurs for a density that lies above 250. (b) *b*_*R*_ = 2*d*_*R*_: The maximum growth rate occurs at a density of 250. In the interval (0, 500) the function *g*(*R*) is symmetric. The shape of this function is very similar to that of the logistic growth function given by eq. (B3) given parameters that fulfill the same constraints (equilibrium at 500 and maximum growth rate of 5). (c) *b*_*R*_ = 4*d*_*R*_: The maximum growth rate occurs for a density that lies below 250. (d) *b*_*R*_ = 10*d*_*R*_: The maximum growth rate is shifted to an even lower density. The consequences of these different growth function for the distribution of bird densities are shown in fig. B1.

#### Maturation dynamics

Logistic growth describes a population of identical individuals. As an alternative, we now derive a growth function when the prey population is stage-structured, with birds feeding on adults that result from maturation of juveniles. We assume that juveniles, with density *J*(*z*) at distance *z*, are produced by adults, with density *R*(*z*) at distance *z*, at a per-capita birth rate *b*_*R*_ and die at a per-capita death rate *d*_*J*_. Juveniles mature into adults at a density-dependent per-capita maturation rate *m*_*J*_/(*k*_*J*_ + *J*(*z*)), where *m*_*J*_ denotes the maximum maturation rate and *k*_*J*_ the density of juveniles at which maturation is half-maximal. Adults move at rate *m*, while juveniles are assumed to be too small to show significant movement. Finally, adults die from causes other than predation by birds at a per-capita death rate *d*_*R*_. Then, at equilibrium the density of juvenile and adult prey fulfill

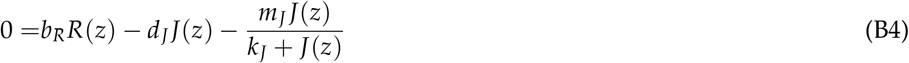

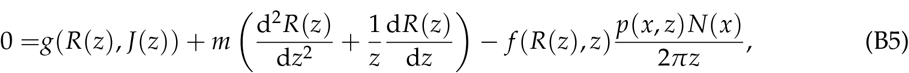

where

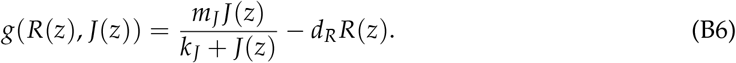

Solving eq. (B4), a quadratic function in *J*(*z*), we find that the relevant root for *J*(*z*) is

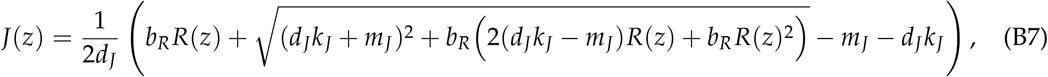

which, on substituting into eq. (B6), results in the growth rate of adult prey taking the form

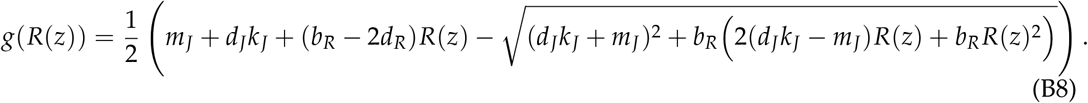

We refer to this growth function as the maturation model. In short, this function describes the growth rate of an adult prey population at distance *z* at equilibrium as it results from maturation of juveniles.

Now recall (eq. B1) that bird density *n*(*z*) is the product of prey renewal *G*(*R*(*z*), *z*) and the net energy value *B*(*R*(*z*), *z*) per prey item. These quantities are shown in fig. B1 as a function of *z* for three different values of the search costs *c*_s_. Note that fig. B1 assumes *m* = 0 so that *G*(*R*(*z*), *z*) = *g*(*R*(*z*), *z*) (see eq. 6). To understand the shape of the renewal function, shown in the fourth row of fig. B1), we start by investigating it as a function of *R*. This is shown in fig. B2 for four different parameter combinations. Using symbolic calculation, it can be shown that eq. (B8) is an unimodal function of *R* that passes through zero at *R* = 0 and *R* = (*m*_*J*_(*b*_*R*_ − *d*_*R*_) − *d*_*J*_*d*_*R*_*k*_*J*_)/(*d*_*R*_(*b*_*R*_ − *d*_*R*_)), which is the equilibrium density for adult prey in the absence of predation by birds. Furthermore, for *b*_*R*_ = 2*d*_*R*_ (fig. B2b), this function has its maximum at half the equilibrium density. Numerical calculations show that the position of the maximum increases if *b*_*R*_ is decreased relative to *d*_*R*_ (fig. B2a) and decreases if *b*_*R*_ is increased relative to *d*_*R*_ (fig. B2c,d). The reason that *g*(*R*) decreases when *R* passes a certain threshold is that the prey population becomes maturation limited– more adults producing offspring results in stronger density dependence among juveniles, and this slows down maturation. This effect becomes more pronounced as *b*_*R*_ increases relative to *d*_*R*_. Hence, the larger *b*_*R*_ relative to *d*_*R*_, the larger the interval of *R*-values for which *g*(*R*(*z*)) decreases. This observation translates to the shape of *g*(*R*(*z*)) when plotted as a function of distance *z*. Panels (c) and (d) in the fourth row of fig. B1, corresponding to the leftward-shifts of the maximum growth rate (due to *b*_*R*_ > 2*d*_*R*_) shown in panels (c) and (d) of fig. B2, show a monotonic decline in the growth rate, simply because the prey equilibrium density at the island, *R*(0), shown in the top row of fig. B1, exceeds the *R*-value where the growth function has its maximum. In contrast, for *b*_*R*_ ≤ 2*d*_*R*_ (column (a) and (b) in fig. B1), whether the function *g*(*R*(*z*)) is unimodal in *z* depends on the search costs *c*_s_. Lower search costs result in a more pronounced prey depletion halo (lower value of *R*(0), see first row in fig. B1), and *g*(*R*(*z*)) is hump-shaped (blue and orange line in fig. B1(a) and blue line in (b)) if the value of *R*(0) is lower than the value of *R* where *g*(*R*) has a maximum (fig. B2) and monotonically decreasing otherwise.

The shape of the graphs for *n*(*z*) (third row in fig. B1) closely follows those for *g*(*R*(*z*)), and are only modified slightly by the change of the net energy value *B*(*R*(*z*), *z*) (fifth row in fig. B1) with distance from the island. In the absence of search costs (*c*_s_ = 0, blue lines), *B*(*R*(*z*), *z*) decreases with *z* due to increasing flight costs. If *c*_s_ is sufficiently high, however, *B*(*R*(*z*), *z*) increases with distance. This is because the time to catch a prey item decreases with increased prey density, so that the costs spend on catching prey decrease with *z*.

The logistic growth function described by eq. B3 is very similar to the curve shown in fig. B2(b), as both curves are symmetric around the *R*-value that is equal to half the equilibrium density in the absence of predation. As a consequence, the graphs in column (b) in fig. B1 are virtually indistinguishable from those that would result from logistic growth. Hence, qualitatively logistic growth can be interpreted as a special case of the maturation model that results when *b*_*R*_ = 2*d*_*R*_.

#### Logistic growth with productivity gradient

Eq. (B3) assumes that the parameters *r* and *α* do not depend on the distance from the island. This assumption approximates the situation around isolated volcanic islands, where the sea floor drops sharply with increasing distance from the island so that the surrounding waters can be considered approximately homogeneous. This assumption is less appropriate for colonies where the surrounding sea shows a gradient in primary productivity with increasing *z*. Such productivity gradients can easily be incorporated in our model. To do this, we extend the logistic growth model (eq. B3) to

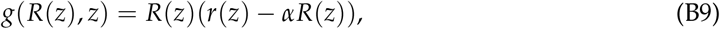

where

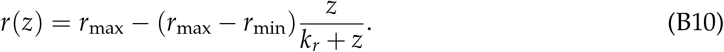

This function for *r*(*z*) is monotonically decreasing in *z*, starting at *r*_max_ for *z* = 0 and approaching *r*_min_ as *z* goes to infinity. The parameter *k*_*r*_ denotes the distance at which the intrinsic growth rate has dropped by 1/2(*r*_max_ − *r*_min_). In the absence of prey movement (*m* = 0), eqs. (11)–(13) apply unchanged.

In eq. (B9), the growth function depends directly on *z*. This generalization of the model does affect the equilibrium prey density and the ideal free distribution, as given by eqs. (11) and (12), respectively, since these results do not hinge on the functional form of *G*(*R*(*z*), *z*). But we need to recalculate the maximum flying distance. To do this, we replace *R*^*^ with *R*^*^ = *r*(*z*_max_)/*α* in eq. (9). With this change, eq. (9) becomes a quadratic equation in *z*_max_, and we find that the relevant root equals

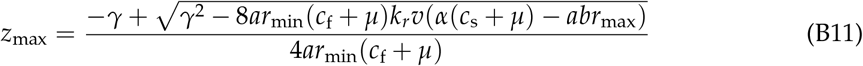

with

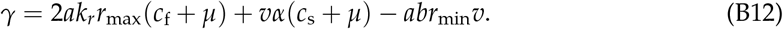

Figure B3 shows three examples of how productivity gradients can affect the distribution of bird densities. The panels in the first row show, for *m* = 0 such that *G*(*R*(*z*), *z*) = *g*(*R*(*z*), *z*), the equilibrium prey density *r*(*z*)/*α* at distance *z* for productivity gradients of increasing strength. Productivity gradients cause prey renewal *g*(*R*(*z*), *z*) (fifth row) to drop sharply close to the island. Depending on the strength of the productivity gradient, this decrease continues until the maximum flying distance (green line in column (b), and all lines in column (c)) or peaks at an intermediate distance. Note, that *g*(*R*(*z*), *z*) also varies with the search costs *c*_s_, as this affects prey abundance *R*(*z*). The peak at intermediate distances under a weak productivity gradient (and under low search costs under an intermediate productivity gradient) results from *g*(*R*(*z*), *z*), when viewed as a function of *R*, having a maximum at an intermediate prey density, just as under the maturation and logistic growth models. The shape of the graphs for *n*(*z*) (fourth row) again closely follows those for *g*(*R*(*z*), *z*), and is only modified slightly by the slope of the net energy content *B*(*R*(*z*), *z*) (sixth row) per prey, just as under the maturation and logistic prey models.

**Figure B3:**
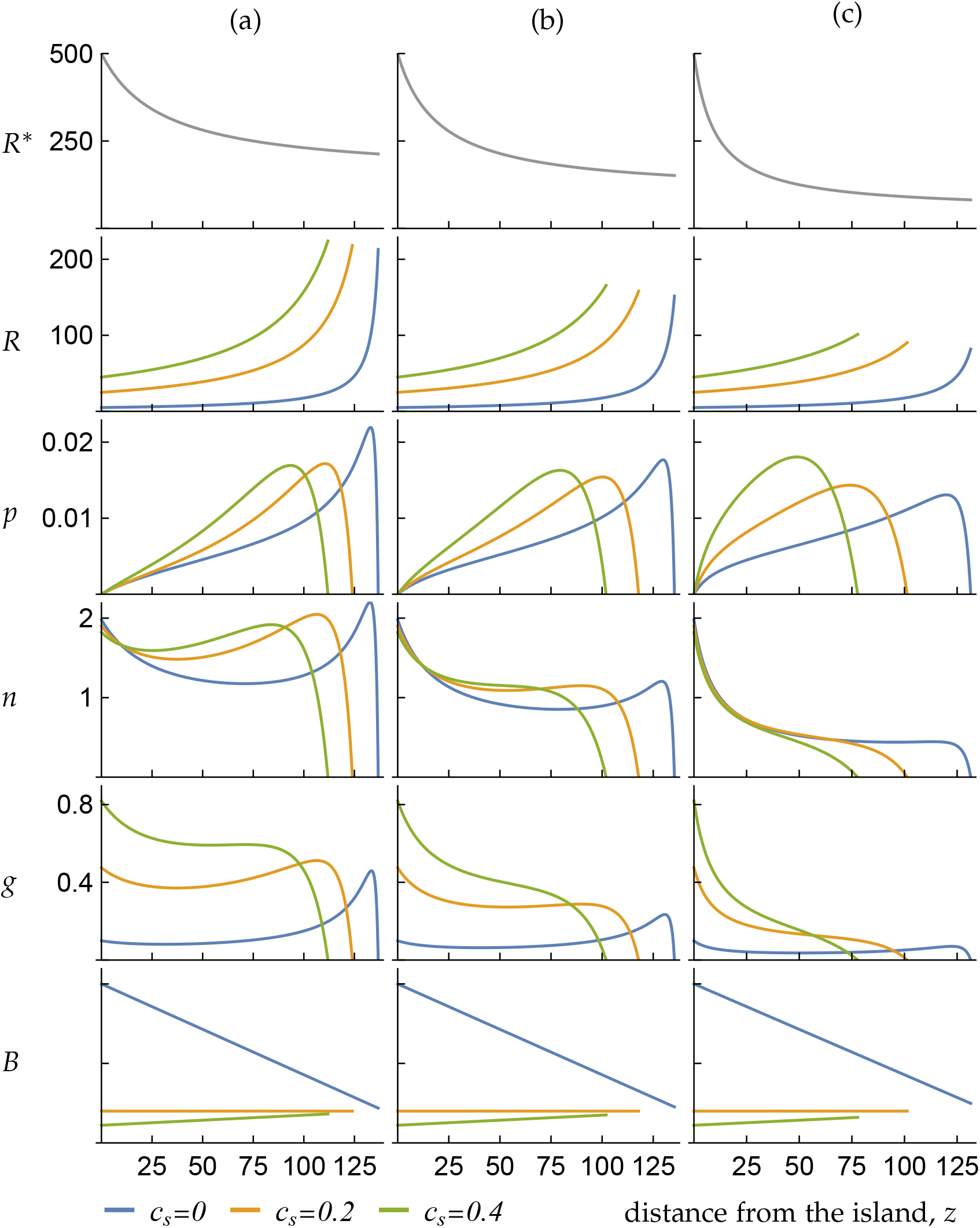
The effect of productivity gradients is illustrated under logistic prey dynamics for three parameter combinations determining the intrinsic growth rate *r*(*z*) as given by eq. (B10) (in columns, parameters: *r*_max_ = 0.02 and (a) *r*_min_ = 0.3*r*_max_, *k*_*r*_ = 30, (b) *r*_min_ = 0.2*r*_max_, *k*_*r*_ = 20, and (c) *r*_min_ = 0.1*r*_max_, *k*_*r*_ = 10) and three values of the search costs *c*_s_ (colored lines). The top row shows the equilibrium prey density *R*^*^ = *r*(*z*)/*α* in the absence of predation by birds, showing that the prey productivity gradient becomes steeper and more pronounced from left to right. The second row shows the equilibrium prey density *R*(*z*) in the presence of predation by birds, that is, the prey depletion halo. Curves for *R*(*z*) of the same color are identical across the three columns but are cut off at different distances, since the maximum flying distance *z*_max_ (eq. B11) decreases with the severity of the productivity gradient. The third row shows the birds’ ideal free distribution *p*(*z*). The fourth row shows the density of birds *n*(*z*) foraging at distance *z*. The pattern shown in the fourth row can be understood based on the fifth (showing *g*(*R*(*z*), *z*)) and sixth (showing *B*(*R*(*z*), *z*)) row. Curves for *B*(*R*(*z*), *z*) of the same color are identical across the three columns but are cut off at different distances, as for *R*(*z*). Parameter values as in fig. 1 but for *m* = 0 and *c*_s_ as indicated by the color code.

#### Conclusion

Our exploration of the effect of three different functions for the growth rate *g*(*R*(*z*)) on the density of birds *n*(*z*) foraging at distance *z*, shows that *n*(*z*) is highly sensitive to details of the prey growth rate. The monotonic and initially steep decline of bird density with increasing distance from the island found by Weber et al. (2021, their fig.1A) is predicted by our model only under sufficiently strong productivity gradients (fig. B3c). In the absence of a productivity gradient, a monotonic decline is only expected when the prey depletion halo is sufficiently shallow so that the equilibrium prey density at the island is higher than the value where the prey growth rate is maximal. For the growth functions analyzed here, this constellation is more likely under the maturation model than under the logistic model and in the presence of high search costs (fig. Appendix Bc,d). Note that, maybe somewhat surprisingly, the shape of the distribution of bird densities also depends on the birds trait values as it is these that determine the depth of the prey depletion halo.

It is important to point out that Weber et al. (2021), when modeling whether the birds breeding on Ascension Island can cause the observed prey depletion halo, assume that the fish population has a uniform distribution at the beginning of the season and becomes increasingly depleted as the breeding season proceeds (the halo expands), with no regrowth of the fish stock during the season. Our model does not allow for such seasonality, and all our results are derived under the assumption that the system is at equilibrium. To conclude, we show that the distribution of the density of foraging birds can take a variety of forms, including dome-shaped and monotonically declining, and accurate predictions would require detailed knowledge of empirical details of the prey renewal dynamics.

## Appendix C Multi-species equilibrium and coexistence

In this appendix, we generalize the single species equilibrium conditions given by eqs. (11)–(13) to a community of interacting species.

### Appendix C.1 Equilibrium conditions for prey, bird behavior and demography for *n* coexisting bird species

According to our assumptions, the equilibrium prey density *R*(*z*), the ideal free distribution *p*(*x, z*) of a species with trait value *x*, and the equilibrium abundance *N*(*x*) of birds with that trait value in a community *χ* = {*x*_1_, *x*_2_, …, *x*_*n*_} are determined by the following coupled system of equations, which generalizes the equilibrium of the single species model of the main text (section 2.2) to *n* species.

First, the prey at distance *z* at equilibrium balances renewal and consumption,

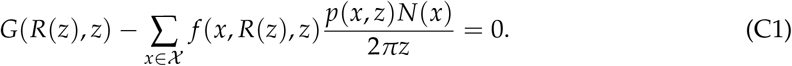

Here, the renewal function *G*(*R*(*z*), *z*) is given by eq. (A10) while prey consumption is obtained from the assumption that the *N*(*x*) birds of the species with trait value *x* are distributed according to the ideal free distribution *p*(*x, z*), and that at distance *z* the *p*(*x, z*)*N*(*x*) birds are homogeneously distributed over the circumference of the circle centered at the island with radius *z*.

Second, the ideal free distribution *p*(*x, z*) for each *x* ∈ *X* is characterized by the fact that the payoff obtained by individuals with trait value *x* ∈ *X* is identical at all distances where these individuals forage. Furthermore, the payoff at distances where individuals forage is at least as high as the payoff that would be obtained at distances where individuals do not forage. Thus, for each *x* ∈ *χ* we have

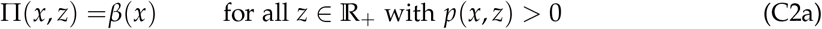

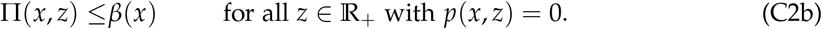

Here, *β*(*x*) is the constant payoff obtained at visited foraging distances to an individual with trait value *x* (the set of foraging distances satisfying eq. (C2a) is the support of the ideal free distribution). While the payoffs given by eq. (C2) do not depend explicitly on *p*(*x, z*) (just as in the single species case), the dependence enters through eq. (C1) as the payoff depends on the equilibrium prey density *R*(*z*), which depends on the ideal free distribution. At the ideal free distribution of the multispecies community characterized by eq. (C2), for each individual with trait value *x* and each *x* ∈ *χ*, we have

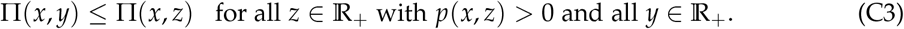

This implies that no individual of any species has an incentive to unilaterally change behavior given all other individuals behave according to the ideal free distribution of their species. Thus, the multi-species ideal free distribution is a multi-species Nash equilibrium.

Third, the species with trait value *x* ∈ *χ* is at its demographic equilibrium *N*(*x*) when birth balances deaths,

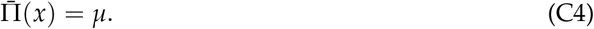

Here,

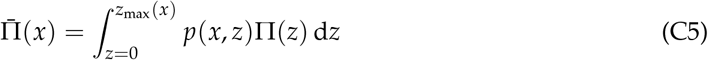

is the mean payoff, which we equate to mean fecundity of an individual with trait value *x*. Furthermore, *z*_max_(*x*) is the maximum traveling distance of such an individual, as given by eq. (10). Here too, the dependence of the birth rate, given by average payoff, depends on *N*(*x*) indirectly through the dependence of the equilibrium prey density on *N*(*x*) (eq. C1).

The coupled system of equations (C1)-(C4) characterizes the prey equilibrium *R*(*z*), the behavioral equilibrium *p*(*x, z*) and the equilibrium bird abundance *N*(*x*).

**Figure C1:**
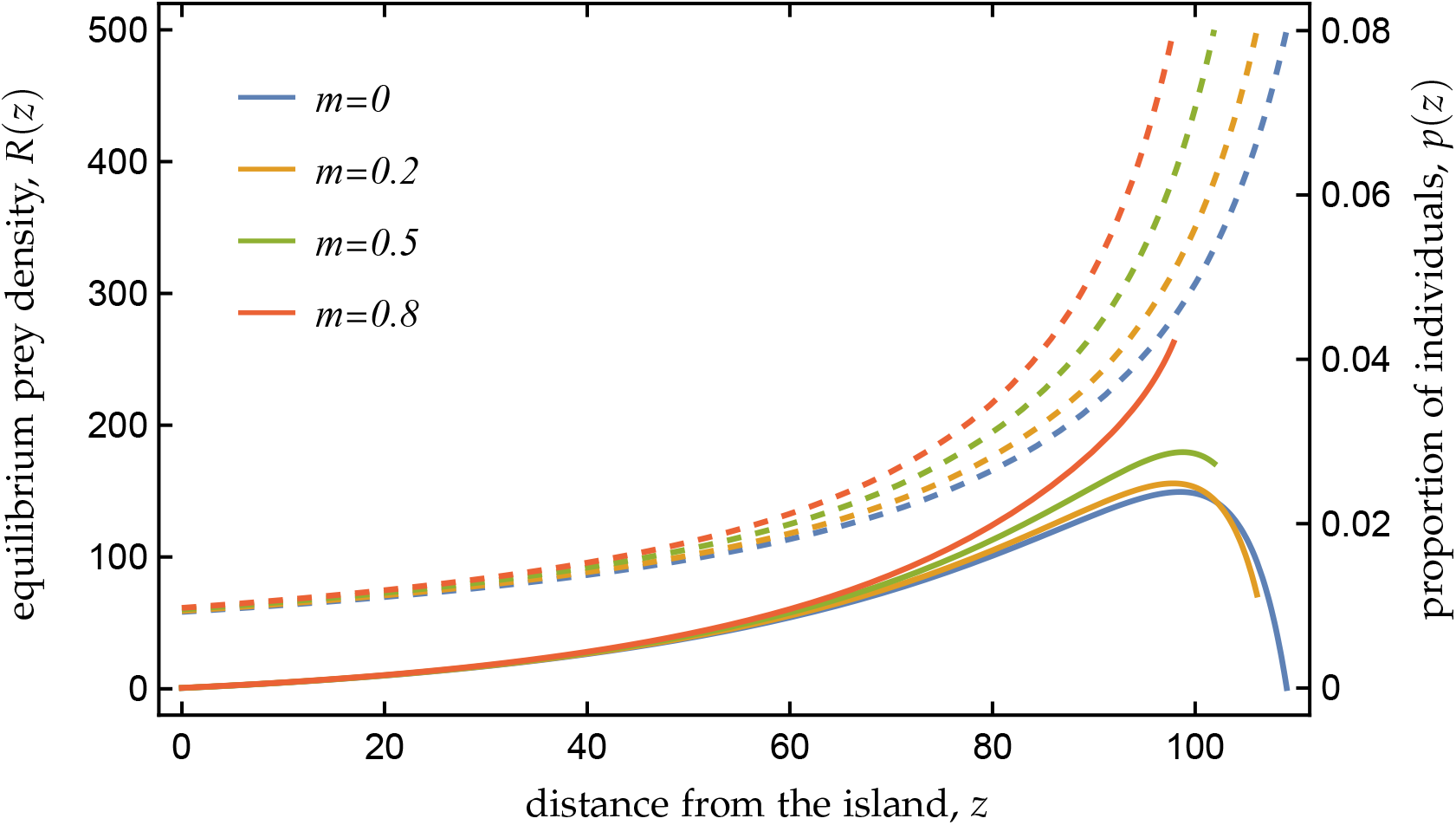
Equilibrium prey density *R*(*z*) (hatched lines, left y-axis) and ideal free distribution *p*(*z*) (proportion of birds feeding at distance *z*, solid lines, right y-axes) as function of distance *z* from the island for five different movement rates *m* and a fixed bird population size of 200000 individuals under logistic prey dynamics. In contrast to the results of bird populations at demographic equilibrium, as shown in fig. 1(a), the maximum flying distance decreases with increasing *m* and prey halos become less pronounced. Parameter values as in fig. 1.

### Appendix C.2 Properties of equilibrium prey densities

To determine the prey equilibrium and the behavioral and demographic equilibrium for multiple bird species, we start by presenting two properties of the equilibrium prey density distribution

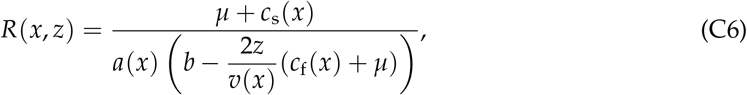

which denotes the equilibrium prey density under the assumption that the bird population is monomorphic for trait value *x* (i.e., eq. 11).

First, for two species with trait values *x*_1_≠*x*_2_ the graphs of the equilibrium prey densities *R*(*x*_1_, *z*) and *R*(*x*_2_, *z*) intersect at most once. To see this, set

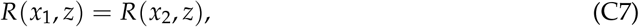

which, owing to eq. (C6), is given by

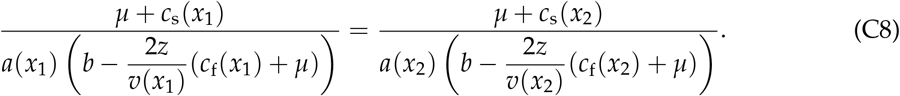

This equality defines a linear equation in *z*, which can have at most one solution. The value of *z* satisfying the equality is the intersection point of the two equilibrium prey density distributions, which may lie beyond *z*_max_(*x*_1_) or *z*_max_(*x*_2_).

Second, *R*(*x, z*) is convex in *z* in the interval [0, *z*_max_(*x*)). This can be seen from the second derivative of eq. (11) with respect to *z*, which is given by

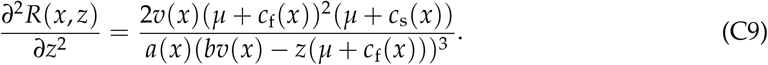

The right-hand side is positive for *z* < *bv*(*x*)/(*µ* + *c*_f_(*x*)). This is true for *z* < *z*_max_ as long as *z*_max_(*x*) < *bv*(*x*)/(*µ* + *c*_f_(*x*)) holds, which is indeed the case as the following shows. By writing eq. (10) as

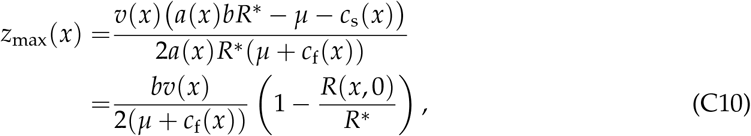

where

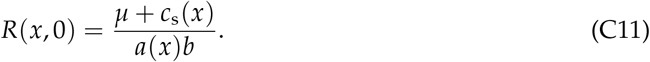

It follows that *z*_max_(*x*) < *bv*(*x*)/(*µ* + *c*_f_(*x*)), because the term in parentheses on the right-hand side of eq. (C10) is less than 1. Thus, *∂*^2^*R*(*x, z*)/*∂z*^2^ > 0 and the function *R*(*x, z*) is convex in *z* for *z* < *z*_max_(*x*).

### Appendix C.3 Mutual exclusive foraging and prey depletion halo

We have now all concepts in place to determine the joint equilibrium conditions for prey, bird behavior and demography in a community of species with trait values *X*. Let us first focus on two species, say species 1 with trait value *x*_1_ and species 2 with trait value *x*_2_. Assume that *R*(*x*_1_, 0) < *R*(*x*_2_, 0) and *z*_max_(*x*_1_) < *z*_max_(*x*_2_) (as an example, consider *x*_1_ = 3.5 and *x*_2_ = 2 with the parameters as in fig. C2). Then, given that the curves *R*(*x*_1_, *z*) and *R*(*x*_2_, *z*) are convex, the above ordering implies that they intersect exactly once, say, at the distance *z* = *z*_1_. Given the assumption *R*(*x*_1_, 0) < *R*(*x*_2_, 0), species 1 depletes prey to a lower density in the interval [0, *z*_1_] and thereby competitively excludes species 2. By contrast, species 2 depletes prey to a lower density in the interval [*z*_1_, *z*_max_(*x*_2_)] and thereby competitively excludes species 1. In conclusion, the interval [0, *z*_max_(*x*_2_)] of utilized foraging distances can be partitioned into two mutually exclusive subintervals, such that [0, *z*_max_(*x*_2_)] = [0, *z*_1_] ∪ [*z*_1_, *z*_max_(*x*_2_)] and *z*_1_ = [0, *z*_1_] ∩ [*z*_1_, *z*_max_(*x*_2_)] (thus, with mutually exclusive we mean that two intervals intersect at most at their boundary). If, however, *R*(*x*_1_, 0) < *R*(*x*_2_, 0) and *z*_max_(*x*_1_) > *z*_max_(*x*_2_), then species 1 excludes species 2 over the whole interval [0, *z*_max_(*x*_1_)] of utilized foraging distances (as is the case for *x*_1_ = 0.5 and *x*_2_ = 0.3 in fig. C2). In summary, starting with a community *χ*= {*x*_1_, *x*_2_}, either the two species coexist by occupying two mutually exclusive distance intervals, or they mutually exclude each other with one species going extinct.

Suppose now that two species coexist and that we add a third species with trait value *x*_3_ (*χ* = {*x*_1_, *x*_2_, *x*_3_}) and that, in addition to the above ordering *R*(*x*_1_, 0) < *R*(*x*_2_, 0) and *z*_max_(*x*_1_) < *z*_max_(*x*_2_), we have, say, *R*(*x*_2_, 0) < *R*(*x*_3_, 0) and *z*_max_(*x*_2_) < *z*_max_(*x*_3_) (as an example, consider *x*_1_ = 3.5, *x*_2_ = 2 and *x*_3_ = 1 with the parameters as in fig. C2). Since “is greater than” (>) is a transitive relation on the real numbers, we then also have *R*(*x*_1_, 0) < *R*(*x*_3_, 0) and *z*_max_(*x*_1_) < *z*_max_(*x*_3_). Due to the convexity of *R*(*x*_1_, *z*), *R*(*x*_2_, *z*) and *R*(*x*_3_, *z*), it follows that (i) the curves *R*(*x*_2_, *z*) and *R*(*x*_3_, *z*) intersect exactly once, say at *z*_2_, and (ii) the intersection points *z*_1_ (of *R*(*x*_1_, *z*) and *R*(*x*_2_, *z*)) and *z*_2_ have the ordering *z*_1_ < *z*_2_. Thus, the interval of utilized foraging distances can now be partitioned into three mutually exclusive distance intervals [0, *z*_max_(*x*_3_)] = [0, *z*_1_] ∪ [*z*_1_, *z*_2_] ∪ [*z*_2_, *z*_max_(*x*_3_)]. In each of these intervals, a different species induces the lowest equilibrium prey density. Hence, starting with a community *χ* = {*x*_1_, *x*_2_, *x*_3_}, either the three species coexist by occupying three mutually exclusive bands (fig. C2 for *x*_1_ = 3.5, *x*_2_ = 2 and *x*_3_ = 1), or two species coexist on mutually exclusive bands (fig. C2 for *x*_1_ = 3.5, *x*_2_ = 2 and *x*_3_ = 0.5), or only one species persists (fig. C2 for *x*_1_ = 1, *x*_2_ = 0.5 and *x*_3_ = 0.3).

By induction, this argument generalizes to a community *χ* = {*x*_1_, *x*_2_, …, *x*_*n*_}. Suppose, the ordering of the maximum flying distances of the *n* species, say, *z*_max_(*x*_1_) < *z*_max_(*x*_2_) < … < *z*_max_(*x*_*n*_) matches the ordering of their equilibrium prey density distributions at distance *z* = 0, *R*(*x*_1_, 0) < *R*(*x*_2_, 0) < … < *R*(*x*_*n*_, 0) (i.e., a larger maximum flying distance implies a higher equilibrium prey density at distance *z* = 0). Then, the interval (0, *z*_max_(*x*_*n*_)) of utilized foraging distances can be partitioned into the *n* distance intervals [0, *z*_1_], [*z*_*i*−1_, *z*_*i*_] for *i* ∈ {2, …, *n* − 1} and [*z*_*n*−1_, *z*_max_(*x*_*n*_)], where *z*_*i*_ (*i* ∈ {1, …, *n* − 1}) is obtained by solving the linear equation

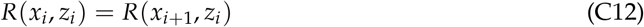

for *z*_*i*_ using eq. (C6). These intervals fulfill

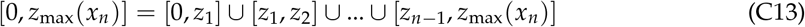

and any two intervals intersect at most at a common boundary. Thus, they are mutually exclusive.

**Figure C2:**
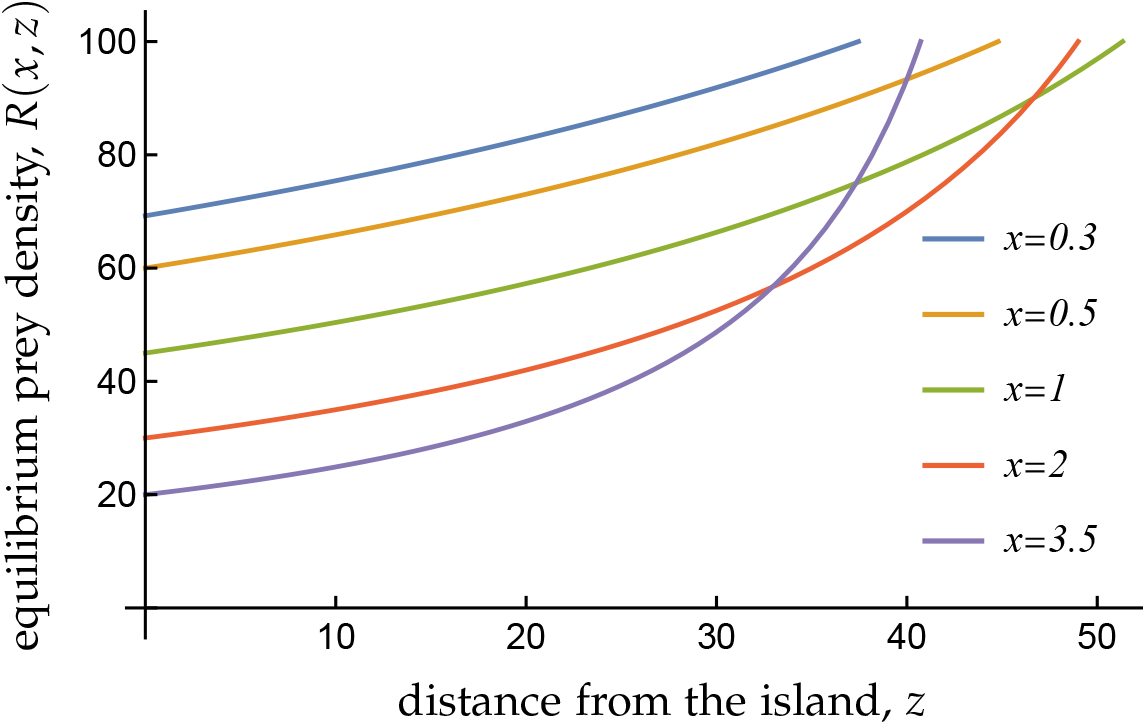
Equilibrium prey density *R*(*x, z*) as a function of distance *z* for five different species, assuming that capture efficiency *a* and flying speed *v* are determined by the trait value *x*. This illustrates that, for certain parameters, not all equilibrium prey density curves intersect at *z*-values less than *z*_max_. The following sequence of community assembly could be envisaged. A community consisting of a species with *x* = 0.3 is replaced by an immigrant (or mutant) with *x* = 0.5, which in turn is replaced by a new species with *x* = 1. However, a further species with *x* = 2 does not replace the species with *x* = 1 but coexist with it (where the species with *x* = 1 is restricted to a small interval at a large distance from the island. Upon the arrival of a species with *x* = 3.5, a community of three coexisting species has emerged. Parameter values as in fig. 1(a) except for *α* = 0.001. As a result, *R*^*^ = 100, which makes condition (D9), specifying when a trade-off between *v* and *a* results in a trade-off between *R*(*x*, 0) and *z*_max_(*x*), more restrictive.

Each interval is occupied by the species that depletes prey to the lowest equilibrium density in that interval, and the equilibrium prey distribution equals

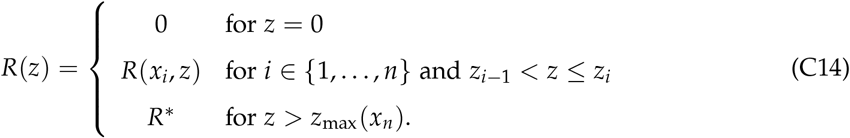

where *z*_*i*−1_ = 0 for *i* = 1, *z*_*i*_ = *z*_max_(*x*_*n*_) for *i* = *n*, otherwise *z*_*i*_ solves eq. (C12). The behavioral strategy equals

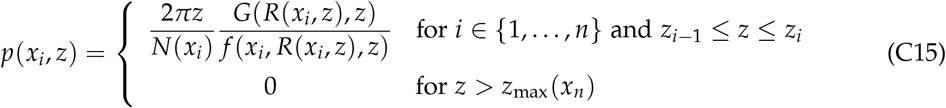

where

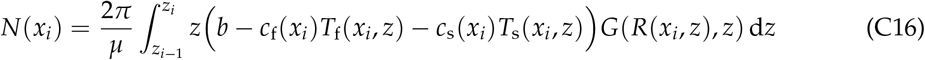

denotes the equilibrium population size of species *i* (compare eq. 13). Eqs. (C14) and (C15) provide an equilibrium prey distribution and ideal free distribution (satisfying 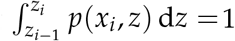), since the system of equations (C1)-(C2) is satisfied. Together, eqs. (C14)-(C16) provide an explicit representation of the joint equilibrium for prey, bird behavior and demography for *n* coexisting bird species. The salient feature is that a single bird species occupies a given interval, and no two species can coexist within the same interval.

## Appendix D Trade-offs

Each species is characterized by a quantitative trait value *x* ∈ R that determines the value of the four foraging components, capture efficiency *a*(*x*), flying speed *v*(*x*), flight costs *c*_f_(*x*), and search costs *c*_s_(*x*). In Appendix C.3, we established that for two species with trait values *x*_1_ and *x*_2_ to be able to coexist due to behavioral niche partitioning, it is necessary and sufficient that the graphs of the equilibrium prey densities – induced by each species in isolation – intersect at some distance *z* that lies within the maximum flying distance of both species. This is the case if and only if *R*(*x*_1_, 0) < *R*(*x*_2_, 0) and *z*_max_(*x*_1_) < *z*_max_(*x*_2_) (or vice versa). In other words, the species that depletes the prey to a lower equilibrium density close to the island also has a lower maximum flying distance. This can be interpreted as a trade-off between a species’ ability to use prey efficiently in the absence of travel costs (close to the island) and in the presence of significant travel costs (resulting in a higher maximum travel distance). In this appendix, we investigate for which pairs of foraging components such a trade-off emerges, which is what we present in table 1 in the main text. We do this in two steps. First, in Appendix D.1 we establish how each foraging component affects *R*(*x*_1_, 0) and *z*_max_(*x*). Second, in Appendix D.2 we ask how changing the trait value *x* affects *R*(*x*_1_, 0) and *z*_max_(*x*). To answer this second questions, we make use of the results from the first step. Assuming that more than two foraging components depend on *x* does not lead to qualitatively new results.

### Appendix D.1 Effects of foraging components on the prey depletion halo

From eqs. (10)–(11) we see that two characteristics of the prey depletion halo, *R*(*x*, 0) and *z*_max_(*x*), can be written explicitly in terms of trait values as

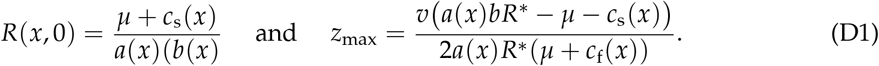

The derivatives of these two quantities with respect to capture efficiency *a*(*x*) are

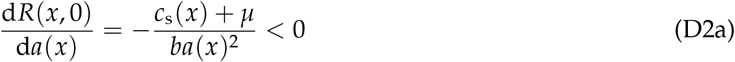

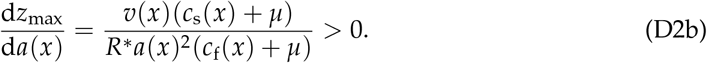

Thus, increasing *a*(*x*) lowers the equilibrium prey density curve *R*(*x, z*) at all distances *z*.The derivatives with respect to flying speed *v*(*x*) are

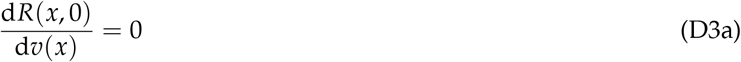

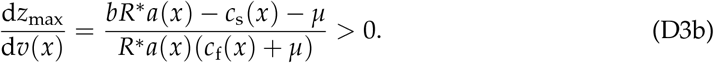

From eqs. (1)-(3) follows that the numerator on the right-hand side of eq. (D3b) can be interpreted as the per-capita growth rate of a species foraging at *z* = 0 and in the absence of competition such that the prey density is given by *R*^*^. For a population to be able to persist, this quantity has to be positive. Thus, increasing *v*(*x*) lowers the equilibrium prey density curve *R*(*x, z*) at all distances *z* > 0.

The derivatives with respect to flying cost *c*_f_(*x*) are

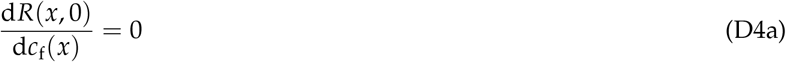

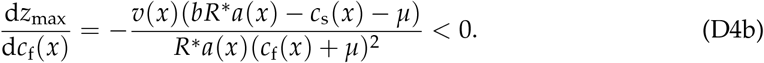

Thus, increasing *c*_f_(*x*) raises the equilibrium prey density curve *R*(*x, z*) at all distances *z* > 0.

Finally, the derivatives with respect to search cost *c*_s_(*x*) are

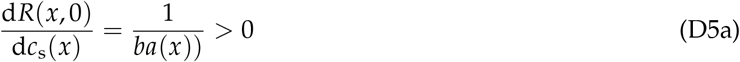

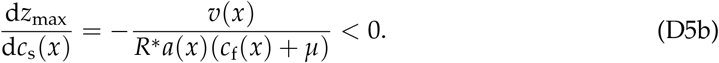

Thus, increasing *c*_f_(*x*) raises the equilibrium prey density curve *R*(*x, z*) at all distances *z*.

### Appendix D.2 Trade-off pairs

In the following calculations, we omit the argument *x* from any foraging component that is not assumed to depend on *x*. Primes denote derivatives with respect to *x*.

For the quantitative trait *x* to have the potential to impose a trade-off between *R*(0, *x*) and *z*_max_(*x*), we assume that *x* maps to the considered foraging components such that the effect of increasing *x* decreases *R*(0, *x*) and *z*_max_(*x*) through one foraging component and increases *R*(0, *x*) and *z*_max_(*x*) through the other foraging component. Changing *x* indeed induces a trade-off if an increase in *x* decreases *R*(0, *x*) and *z*_max_(*x*) (increased competitiveness close to the island at the cost of decreased competitiveness far way from the island), or vice versa.

#### Trade-off between capture efficiency *a*(*x*) and flight speed *v*(*x*)

Based on eqs. (D2) and (D3), to impose a trade-off we assume

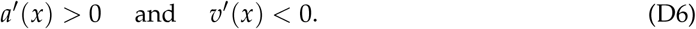

For the derivatives, we find

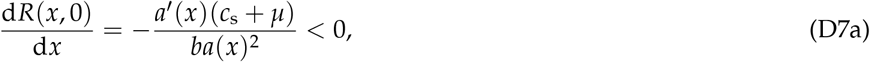

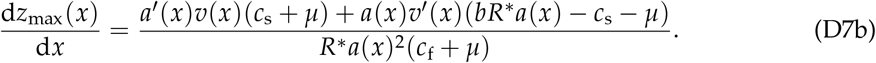

For a trade-off to exist, the right-hand side of eq. (D7b) has to be negative. This is equivalent to

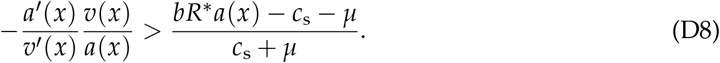

Due to eq. (D6), the function *v*(*x*) and is invertible and differentiable with derivative 1/*v*^*′*^(*x*). With this, we can rewrite the last inequality as

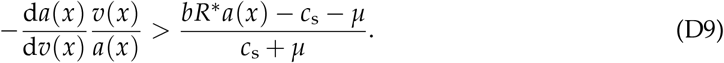

The left-hand side of this inequality gives the slope of the parametric curve that plots *a*(*x*) and *v*(*x*) as a function of *x*, to which we refer as trade-off curve, weighted by the inverse ratio of the two foraging components. Note, that, due to (D6), this trade-off curve has a negative slope. The left-hand side of inequality(D9) is known as the elasticity of the trade-off curve (the percentage change in *a*(*x*) with percentage change in *v*(*x*); e.g., Sydsæter and Hammond, 1995, p. 171-4).

#### Trade-off between capture efficiency *a*(*x*) and flight costs *c*_f_(*x*)

Based on eqs. (D2) and (D4), to impose a trade-off we assume

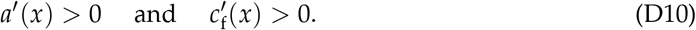

For the derivatives we find

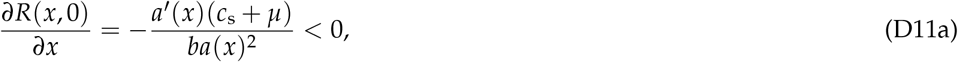

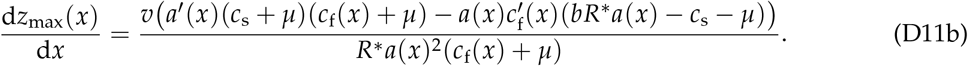

**Figure D1:**
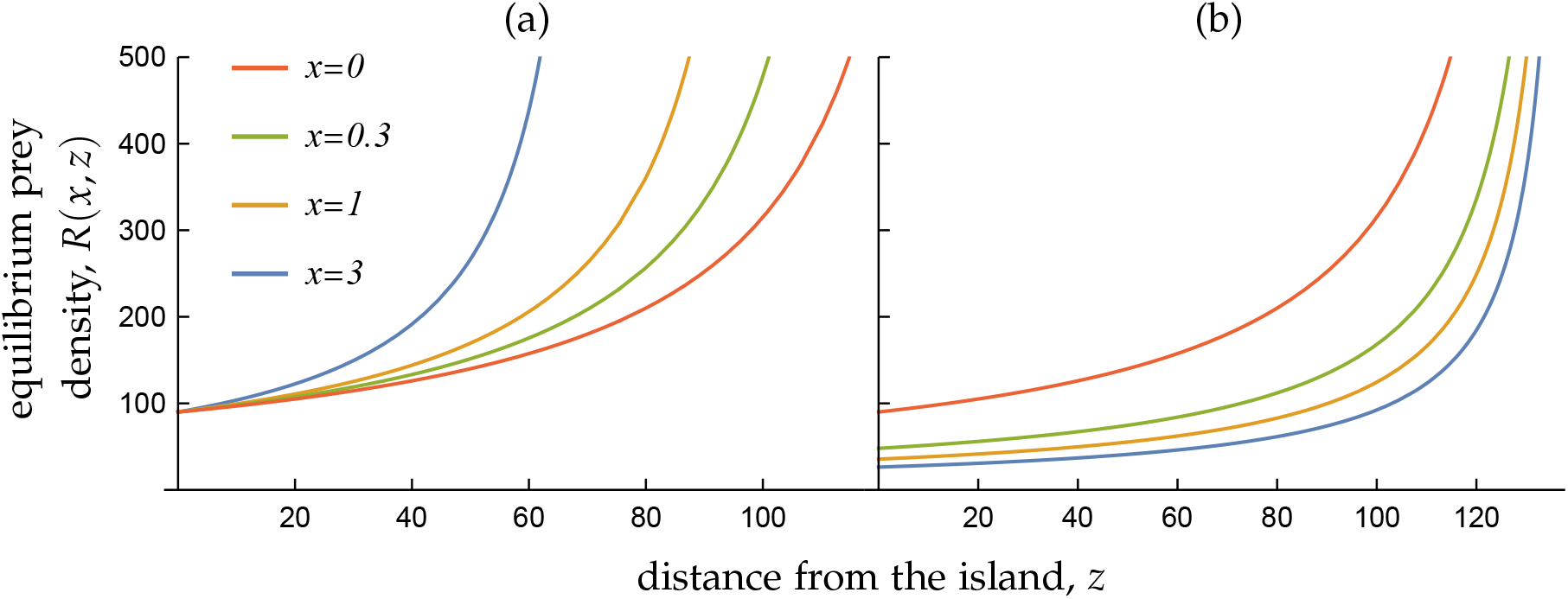
Equilibrium prey density *R*(*x, z*) as a function of distance *z* for four different trait values *x*, assuming that (a) flying speed *v*(*x*) and flight costs *c*_f_(*x*) and (b) capture efficiency *a*(*x*) and search costs *c*_s_(*x*) are trait-dependent. In the first case, all equilibrium prey density curves intersect at *z* = 0 while they never intersect in the second case. In both cases, coexistence of different species is impossible and the species with the lowest equilibrium prey density curve excludes all others (*x* = 0 in (a) and *x* = 10 in (b)). The functions *a*(*x*) and *v*(*x*) are as described in the legend of fig. 2. For flying and search costs, we use *c*_f_(*x*) = *c*_f0_/(1 + *c*_f1_*x*) and *c*_s_(*x*) = *c*_s0_/(1 + *c*_s1_*x*), respectively, with parameters *c*_f0_ = 0.2, *c*_fb_ = 0.01, *c*_s0_ = 0.4 and *c*_s1_ = 0.1. Other parameter values as in fig. 1(a).

For a trade-off to exist, the right-hand side of eq. (D11b) has to be negative. Factoring out 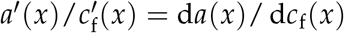, this is equivalent to

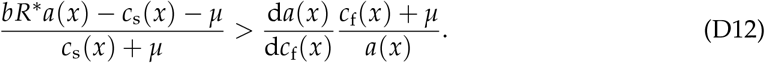

#### Trade-off between capture efficiency *a*(*x*) and search costs *c*_s_(*x*)

In this case, solving eq. (C8) for *z* shows that different equilibrium prey density functions do not intersect with each other (as in fig. D1b). Thus, for a given set of species *𝒳*, the species *x* corresponding to the lowest equilibrium prey density curve excludes all other species and coexistence is impossible.

#### Trade-off between flight speed *v*(*x*) and flight costs *c*_f_(*x*)

In this case, solving eq. (C8) for *z* shows that different equilibrium prey density functions always intersect with each other at *z* = 0 (as in fig. D1a). Thus, for a given set of species *X*, the species *x* corresponding to the lowest equilibrium prey density curve excludes all other species and coexistence is impossible.

#### Trade-off between flight speed *v*(*x*) and search costs *c*_s_(*x*)

Based on eqs. (D2) and (D5), to impose a trade-off we assume

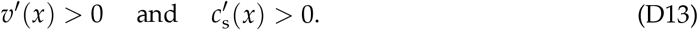

For the derivatives we find

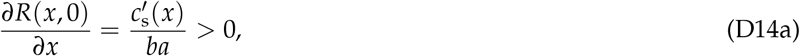

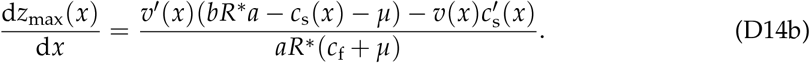

For a trade-off to exist, the right-hand side of eq. (D14b) has to be positive. Factoring out 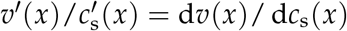, this is equivalent to

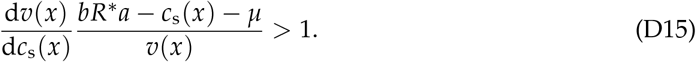

#### Trade-off between flight costs *c*_f_(*x*) and search costs *c*_s_(*x*)

Based on eqs. (D4) and (D5), to impose a trade-off we assume

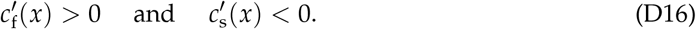

For the derivatives we find

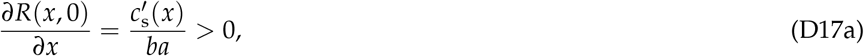

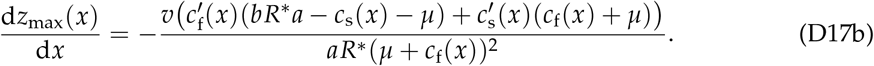

A trade-off exists if eq. (D17b) is positive. Factoring out 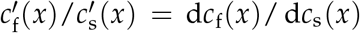, this is equivalent to

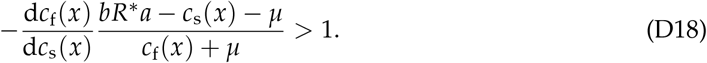

## Appendix E Group foraging

We here derive the model of group foraging described in Box 1. Thus, we keep all assumptions of the foraging process the same as under the model for individual foraging, except that (i) all individuals express an identical foraging behavior, (ii) population (group) size *N* is fixed, and (iii) resources show no movement (*m* = 0). Since we consider groups consisting of a single species, we henceforth omit the argument *x* from all variables.

From the model of individual foraging we know that a maximal foraging distance *z*_max_ exists beyond which payoff is negative and distances beyond this point will never be visited in any behavioral equilibrium. Accordingly, we can write the expected payoff to an individual with behavioral strategy *p* : [0, *z*_max_] → *R*_+_ under group foraging as

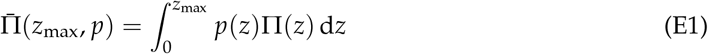

subject to the constraint that

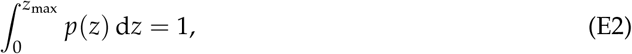

and where

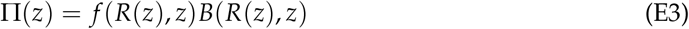

is the payoff from foraging at distance *z*. Eq. (E1) is the net rate of energy delivery of an individual under group foraging and can equivalently be interpreted as the foraging group’s average percapita energy delivery. Here, *B*(*R*(*z*), *z*) and *f* (*R*(*z*), *z*) are defined as in eqs. (1) and (2) in main text. Furthermore, analogous to eq. (5) in the absence of movement (*m* = 0), the equilibrium prey density *R*(*z*) at distance *z* satisfies

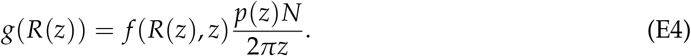

Thus, the mean payoff defined by eq. (E1) is equivalent to that defined by eq. (4).

The behavior *p* in eqs. (E1) and (E4), however, does not yet represent an equilibrium. To find the equilibrium, we have to maximize the payoff 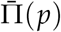 with respect to *p* and the maximum foraging distance *z*_max_ under the integral constraints given by (E2). To solve this optimization problem, we follow standard optimization techniques using calculus of variations (e.g. Bryson and Ho, 1975; Sydsæter et al., 2008). We note that our problem is equivalent to an optimal control problem with a free boundary point and an integral constraint (e.g., Bryson and Ho, 1975, section 2.8 and 3.1). Accordingly, we construct the Lagrangian function

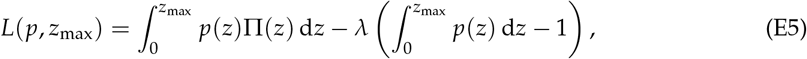

where *λ* is the Lagrange multiplier associated to the integral constraint. The necessary first-order conditions for the function *p* and the distance *z*_max_ to maximize payoff are

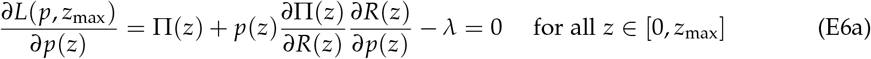

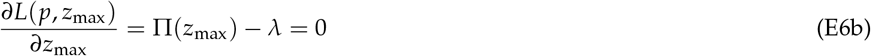

(a special case of Bryson and Ho, 1975, eqs. 2.8.17 and 2.8.20 with a Lagrangian function given by their eq. 3.1.4). Note, that for simplicity of presentation, in the second term on the right-hand side of eq. (E6a) we omit writing the functions explicitly in terms of the arguments with respect to which partial derivatives are taken. The first term in this derivative, Π(*z*), describes the payoff gain to an individual from increasing its tendency to forage at distance *z* and is thus a benefit. The second term describes the change in payoff that results from increased prey depletion at a distance *z* that comes with an increased tendency to forage at that distance. The Lagrange multiplier *λ* can be interpreted as the payoff that is lost at equilibrium if the constraint could be loosened infinitesimally (e.g., Sydsæter et al., 2008). eq. (E6a) says that the gain from foraging at distance *z* must be balanced by the cost of increasing competition from foraging at *z* and the cost this entails. Finally, note that by comparing eq. (E6b) and eq. (E5) it follows that *p*(*z*_max_) = 0.

Eqs. (E6a)–(E6b) show that a candidate equilibrium density *p*(*z*) at *z* has to satisfy

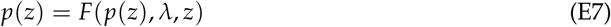

with

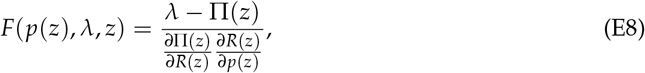

where

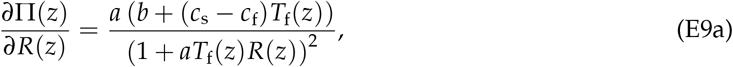

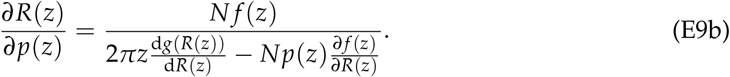

The last equality is obtained by implicit differentiation of eq. (E4) and where *∂ f* (*z*)/*∂R*(*z*) depends on *p*(*z*) owing to eq. (E4). The argument *p*(*z*) in *F*(*p*(*z*), *λ, z*) emphasizes that the density *p*(*z*) in eq. (E7) is defined only implicitly. But this density can be solved by finding the solution *p*(*z*) satisfying eq. (E7) if *λ* is known, which in turn can be obtained by solving

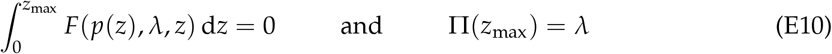

for *λ* and *z*_max_.

In conclusion, once the nature of the prey’s growth rate *g*(*R*(*z*)) is specified, the system of equations given by eqs. (E7)–(E10) in combination with eqs. (E3)–(E4) allows us to compute the candidate behavioral equilibrium (*p, z*_max_), consisting of a foraging distribution and a maximal foraging distance, fulfilling eq. (E7). To ascertain that this is indeed a maximum, one would need to evaluate second-order or sufficiency conditions for a local or global maximum. The literature on control theory does not provide any simple recipe to check any such conditions in the context of free boundary point optimal control problems (e.g., Seierstad and Sydsæter, 1987, Theorem 13, p. 145), and our problem is furthermore compounded with an integral constraint. Hence, we argue somewhat heuristically that the equilibria identified numerically in fig. 3 are indeed maxima. For this, we note that, because there is no prey at *z* = 0 and because payoff decrease with distance *z* in an environment without competitors, the optimal solution has to be a function *p* : [0, *z*_max_] → *R*_+_ with *z*_max_ > 0 and *p*(*z*) > 0 for all *z* ∈ (0, *z*_max_]. Second, in the numerical analysis underlying fig. 3, we find a single positive solution *p*(*z*) for each *z* ∈ (0, *z*_max_] satisfying eq. (E7). This thus has to characterize a maximum.

